# In-situ observation of mitochondrial biogenesis as the early event of apoptosis

**DOI:** 10.1101/2020.08.23.263152

**Authors:** Chang-Sheng Shao, Xiu-Hong Zhou, Yu-Hui Miao, Peng Wang, Qian-Qian Zhang, Qing Huang

## Abstract

Mitochondrial biogenesis is a cell response to external stimuli which is generally believed to suppress apoptosis. However, during the process of apoptosis, whether mitochondrial biogenesis occurs in the early stage of the apoptotic cells remains unclear. To address this question, we constructed the COX8-EGFP-ACTIN-mCherry HeLa cells with recombinant fluorescent proteins respectively tagged on the nucleus and mitochondria, and monitored the mitochondrial changes in living cells exposed to gamma-ray radiation. Besides in situ detection of mitochondrial fluorescence changes, we also examined the cell viability, nuclear DNA damage, reactive oxygen species (ROS), Mitochondrial superoxide, citrate synthase activity, ATP, cytoplasmic and mitochondrial calcium, mitochondrial DNA copy number and expression of transcription genes related to mitochondrial biogenesis as well as the apoptosis biomarkers. As a result, we confirmed that significant mitochondrial biogenesis took place preceding the radiation-induced apoptosis, and the change of mitochondrial biogenesis at early time was closely correlated with the apoptotic cells at late stage. The involved mechanism was also discussed.

**Highlights:**

- A dual fluorescence reporter system was successfully constructed for in-situ observation of mitochondrial biogenesis in living cells.
- The whole process of radiation-induced mitochondrial biogenesis and apoptosis was scrutinized.
- The conception of the relationship between mitochondrial biogenesis and apoptosis was revised.
- Assessment of the early event of mitochondrial biogenesis is critical for prediction of the late fate of cells.

**Graphical abstract:** 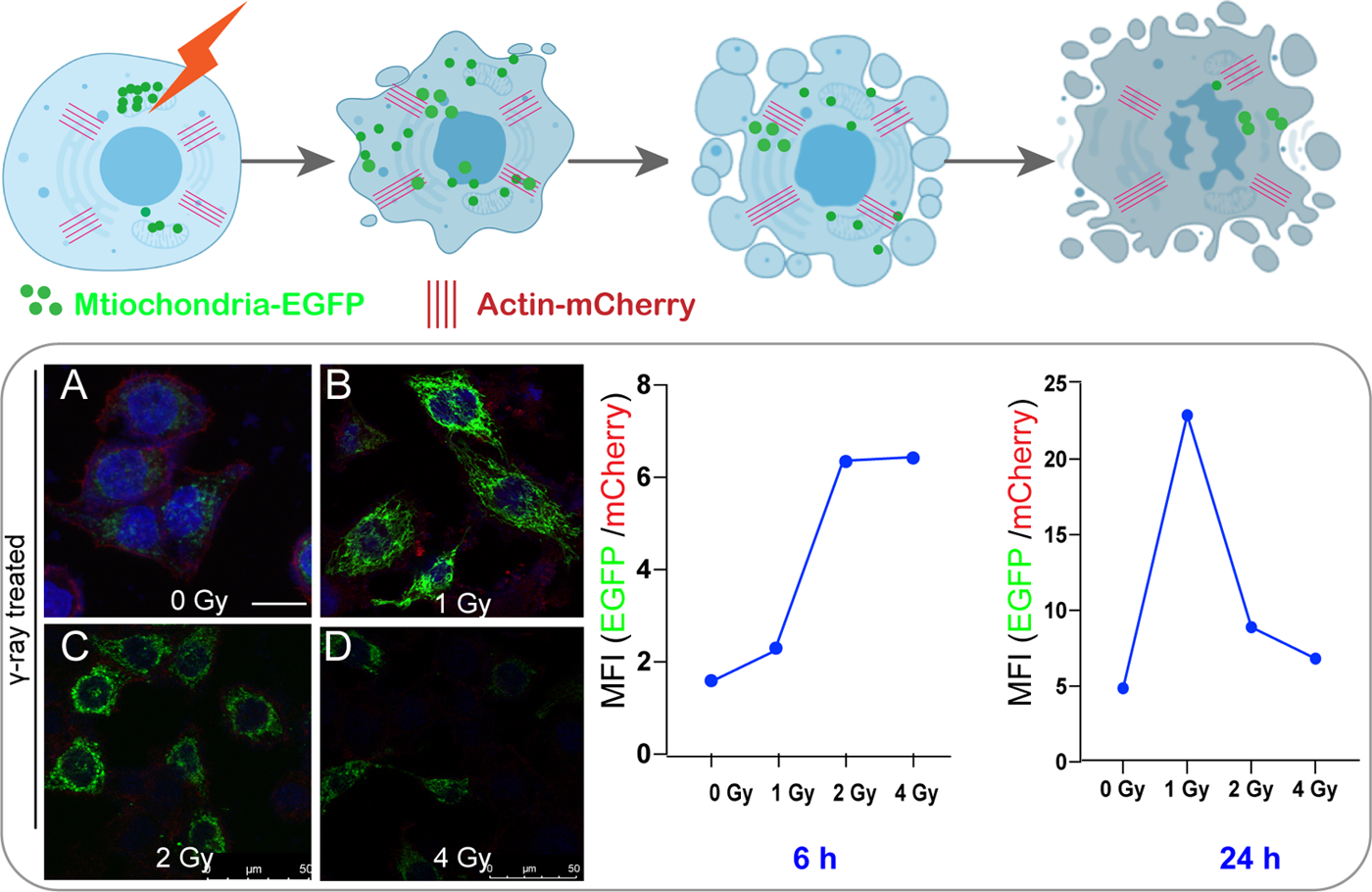

## INTRODUCTION

Mitochondrial biogenesis is a physiological response of cells to external stress that may cause increase of energy demand, and it plays an important role in cell metabolism regulation, signal transduction and mitochondrial ROS regulation (Luo et al., 2016; Vyas et al., 2016). Mitochondrial biogenesis maintains cell homeostasis by ensuring the quality of mtDNA and regulating the renewal of organelles (Yambire et al., 2019), and it has been extensively explored in recent years due to the relevant interests in human aging, neurodegenerative diseases, cell metabolic diseases and tumors(Fanibunda et al., 2019).

Since mitochondrial regulation plays a very critical role in controlling the cell fate, it is also intriguing for researchers to explore the relationship between mitochondrial biogenesis and apoptosis. Generally, apoptosis is mediated by the activation of the caspase pathways which are associated with mitochondrial damage accompanied by destruction of electron transfer, oxidative phosphorylation, ATP production and change of cell redox potential(Burke, 2017). In fact, some studies have claimed that there is a negative correlation between mitochondrial biogenesis and apoptosis, i.e., apoptosis is normally suppressed by mitochondrial biogenesis. For example, it has been reported that Ca^2+^-mediated apoptotisis can be inhibited by enhanced mitochondrial biogenesis (Dam et al., 2013), or, on the other hand, apoptosis may occur by inhibiting mitochondrial biogenesis(Cao et al., 2017). Also, in the study of radiation-induced apoptosis, mitochondrial biogenesis was observed and analyzed (Rai et al., 2018), and it was claimed that enhancement of mitochondrial biogenesis could significantly reduce the proportion of apoptosis caused by ionizing radiation(Yu et al., 2013). However, concerning the whole process of apoptosis, whether and how mitochondrial biogenesis takes place in the early period in the apoptotic cells exposed to the external stimuli such as ionizing radiation remains elusive.

In order to scutinize the relationship between mitochondrial biogenesis and apoptosis, it is thus crucial to monitor the mitochondrial changes in the whole process of apoptotic cells. At present, there are not many methods available to study the influence of external factors on mitochondrial changes. One of the conventional methods is to probe the change of mitochondrial gene copy numbers(Yu, 2011). Another approach is to use fluorescent proteins. Fluorescent proteins are beneficial for studying the living cells with reporting gene expression, which can provide information on protein locations and expression levels as well as the involved biochemical activities(Newman et al., 2011). Actually, study of mitochondrial process using fluorescent proteins is now gaining increasing attention (Katajisto et al., 2015; Melentijevic et al., 2017; Ruan et al., 2017). Recent development is to make use of multiple different fluorescent proteins with different excitation/emission spectra, so that the precision of transcriptional measurements can be ensured by using the reference color for normalization (Miyashiro and Goulian, 2007). In such a circumstance, the ratio of dual fluorescence can be used to eliminate the possible fluorescence measurement errors(Miyashiro and Goulian, 2007). So, with application of this dual fluorescence approach, nowadays researchers are able to observe and analyze the temporal and spatial life trajectories of living cells (Jadhav and Shivdasani, 2019), and maybe more importantly, to scrutinize the early events leading to various bio-effects such as apoptosis as end-point fate in the living cells (Yu et al., 2020).

Therefore, in the present work, we attempted to apply the dual recombinant fluorescent proteins to monitor the mitochondrial biogenesis induced by injuring radiation, and in this way, to examine whether and how the early event of mitochondrial biogenesis would be related to the prominent bio-effects such as apoptosis. For the effect of radiation as the external stimuli to the cells, it is known that ionizing radiation (such as α-, γ-, X-rays, protons, heavy ions) can readily cause various detrimental damages to living cells, which can lead to the change of genetic information, cell mutations, genomic instability, and apoptosis (O’Driscoll and Jeggo, 2006). Mitochondrion, as the important subcellular organelle, is one of the major targets of ionizing radiation(Leach et al., 2001). On the other hand, radiation can also effectively induce mitochondrial biogenesis (Das et al., 2020). For the establishment of the dual fluorescent system for in-situ observation of mitochondrial biogenesis in living cells, we constructed the stable fluorescent reporter cell lines by lentiviral transfection to achieve mitoGFP and nuclear mCherry. As a result, we fulfilled the task of in-situ and real-time observation of mitochondrial changes in the living cells, and observed the effect of mitochondrial biogenesis occurring in the early process of the apoptotic cells. By analyzing the fluorescence ratio of mitoGFP and nuclear mCherry fluorescence, we could therefore more reliably examine the relationship between the early mitochondrial biogenesis and late apoptosis. The detailed mitochondrial/apoptotic processes were analyzed, and the involved mechanism was discussed.

## RESULTS

### Construction of dual-fluorescent stably transfected cell lines

In order to construct a universal dual-fluorescence gene reporter vector, we considered to make use of the simplicity of vector construction and referred to the previously reported dual-fluorescence reporter system, and so we chose the CMV promoter to express the dual-fluorescence target proteins. The mitochondrial reporter expression system is generally localized in the stroma, targeting the leader peptide (MSVLTPLLLRGLTGSARRLPVPRAKIHSLGDP) on the VIIIa subunit of cytochrome C oxidase (COX8) (Haggie and Verkman, 2002) (Figure 1A). As such, we constructed the pLVX-mCherry-actin lentiviral vector (Figure 1B).

**Figure 1.**
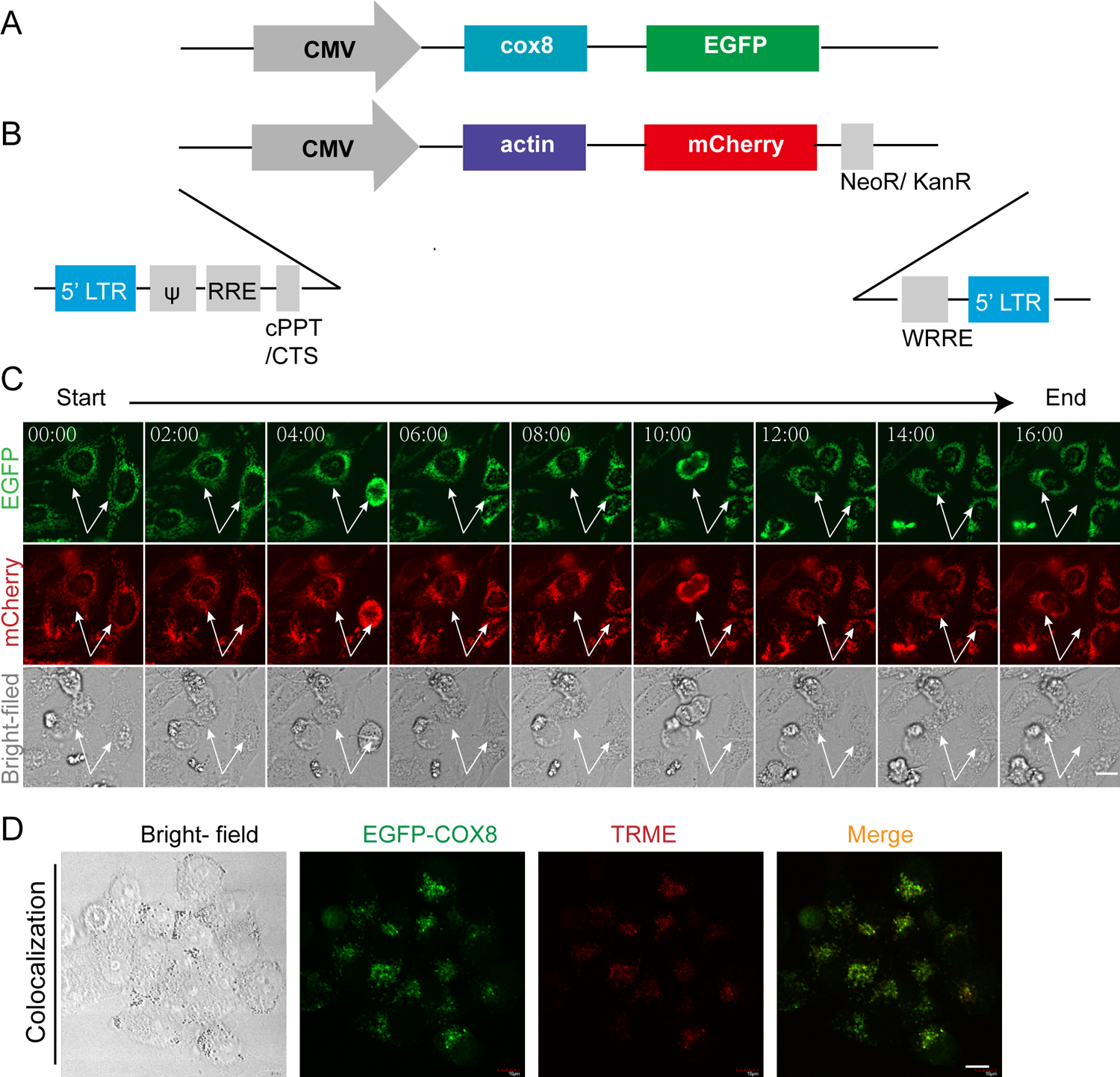
Generation of the dual-fluorescent protein reporter system. (A) Schematic illustration of the pEGFP-cox8 reporter system. (B) Schematic illustration of the pLVX-actin-mCherry reporter system. (C) Live imaging of EGFP-Cox8-mCherry-Actin-HeLa cells within 16 hours. Arrow points to the observed dynamic changes of cell proliferation. (D) Confocal microscopy observation of co-localization of GFP-COX8 on the mitochondria. The GFP-COX8-HeLa cells were stained with TMRE (200 nM) for 20 minutes before the examination by confocal microscopy. Scale bar: 10 μm.

To establish the cell line stably co-expressing COX8-EGFP and ACTIN-mCherry, we transfected the HeLa cells with the plasmid containing the expression sequence of COX8-EGFP, and obtained the COX8-EGFP expression cells. After screening by G418 for 2 weeks, the stably transfected cell line was obtained. The COX8-EGFP-mCherry-ACTIN co-expressing HeLa cell line was further established based on the COX8-EGFP expressing cells with lentiviral transfection system. From the as-constructed HeLa cells, we could monitor the cell activities in real time using the living cell workstation. As shown in Figure 1C, with the change of time, we observed the stable expression of GFP located in mitochondrial cytochrome c oxidase, and also confirmed the stable expression of mCherry associated with the actin proteins in the cells. To be noted, the intensity of mCherry fluorescence associated with actin expression was relatively quite constant with the change of time, indicating that actin fluorescence could be applied as the internal standard for the fluorescence reporting system.

In order to further confirm the GFP expression right on the targeted mitochondria, we then labeled the mitochondria with TMRE dye in the GFP-tagged cells and then examined the dye-stained cells with a confocal microscope. As shown in Figure 1D, the COX8 protein fluorescence (Green) proteins and mitochondrial fluorescence (Red) were indeed co-localized in the cells.

In order to verify the universal validity of our as-constructed dual fluorescence reporting system for observing mitochondrial biogenesis, we also used mitochondrial biogenesis’s initiator 5-Aminoimidazole-4-carboxamide ribotide (AICAR) as for the in-depth testing, or, as the positive control (Komen and Thorburn, 2014). The applied drug concentration was 0.5 mM. As shown in the Figures 2A-B, compared with the non-AICAR treatment group, the AICAR treated group showed increased fluorescence ratio with time. This experiment therefore confirmed that the dual fluorescence system we constructed is feasible and effective for monitoring mitochondrial biogenesis. Furthermore, the correlation between GFP and mitochondrial biogenesis related protein expression were also confirmed by the WB assay (Figures 2C-D). We testified the co-expression levels of GFP protein together with the mitochondrial target protein (e.g. COX8) for the cells treated by AICAR. Technically, cytochrome c (CytoC) was measured in the experiment instead of COX8 as COX8 is the subunit of CytoC oxidase. As shown in Fig. 2C, both GFP and CytoC were in the same trend of expression of levels for the mitochondrial biogenesis proteins including PGC1-α, NRF1 and TFAM.

**Figure 2.**
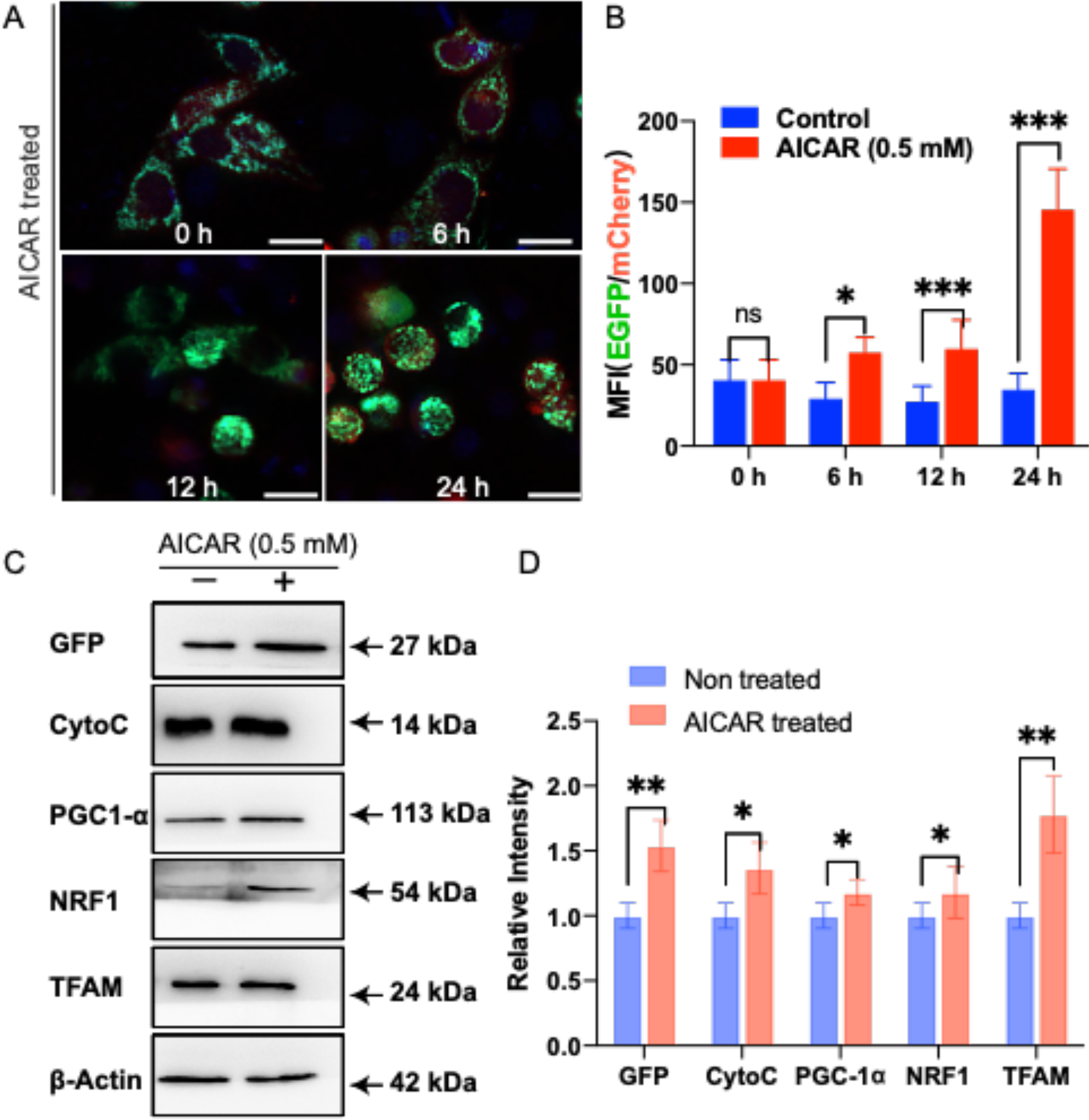
Verification of mitochondrial biogenesis phenomenon observed by the dual fluorescence reporting system (A) The cells were treated with mitochondrial biogenesis inducer drug (AICAR, 0.5 mM) and observed using high-content imaging and screening (HCS) system after different treatment hours. In the figures, green color indicates the mitochondria encoding COX8-EGFP, red color shows the actin encoded by actin-mCherry, and blue color represents the DAPI stained nuclei. Scale bar: 25 μm. (B) The mean fluorescence intensity (MFI) ratios of HeLa cells treated by AICAR after different hours, in comparison with the non-AICAR treated cells as the control group. (C) Western blot (WB) assay for the HeLa cells which were treated with mitochondrial biogenesis inducer drug (AICAR, 0.5 mM). Total proteins were extracted after 12 hours. (D) The graph shows the comparison of the relative intensity for expressions of GFP and mitochondrial biogenesis marker proteins between the non-AICAR treated group and the AICAR treated group. Bar graphs are presented as mean ± sd, n = 3.

Besides, we also examined the AICAR-induced biomarker proteins expression levels related to mitochondrial biogenesis. Figure 2D shows that for the cells treated with AICAR, the expression levels of GFP, CytoC, PGC1-α, NRF1 and TFAM increased significantly in comparison with the untreated group. Also, with the increase of AICAR treatment time, the protein expression levels of mitochondrial biogenesis marker proteins (PGC1, NRF1, TFAM) also increased concomitantly (see Figure S1). All these results therefore confirmed the validity of our fluorescence report system suitable for mitochondrial biogenesis monitoring.

### In-situ observation of the dual fluorescence reporter system for the evaluation of the radiation-induced mitochondrial changes

With the establishment of this dual-fluorescence reporter system, we then employ the laser fluorescence confocal microscope to observe the biological effect in the living cells with the treatment of gamma-ray (γ-ray) irradiation. The COX8-EGFP encoding mitochondria (Green), the ACTIN-mCherry encoding actin (Red), and the DAPI-stained nuclei were observed. The representative images together with the corresponding quantitative analysis of fluorescence intensity are shown in Figure 3A.

**Figure 3.**
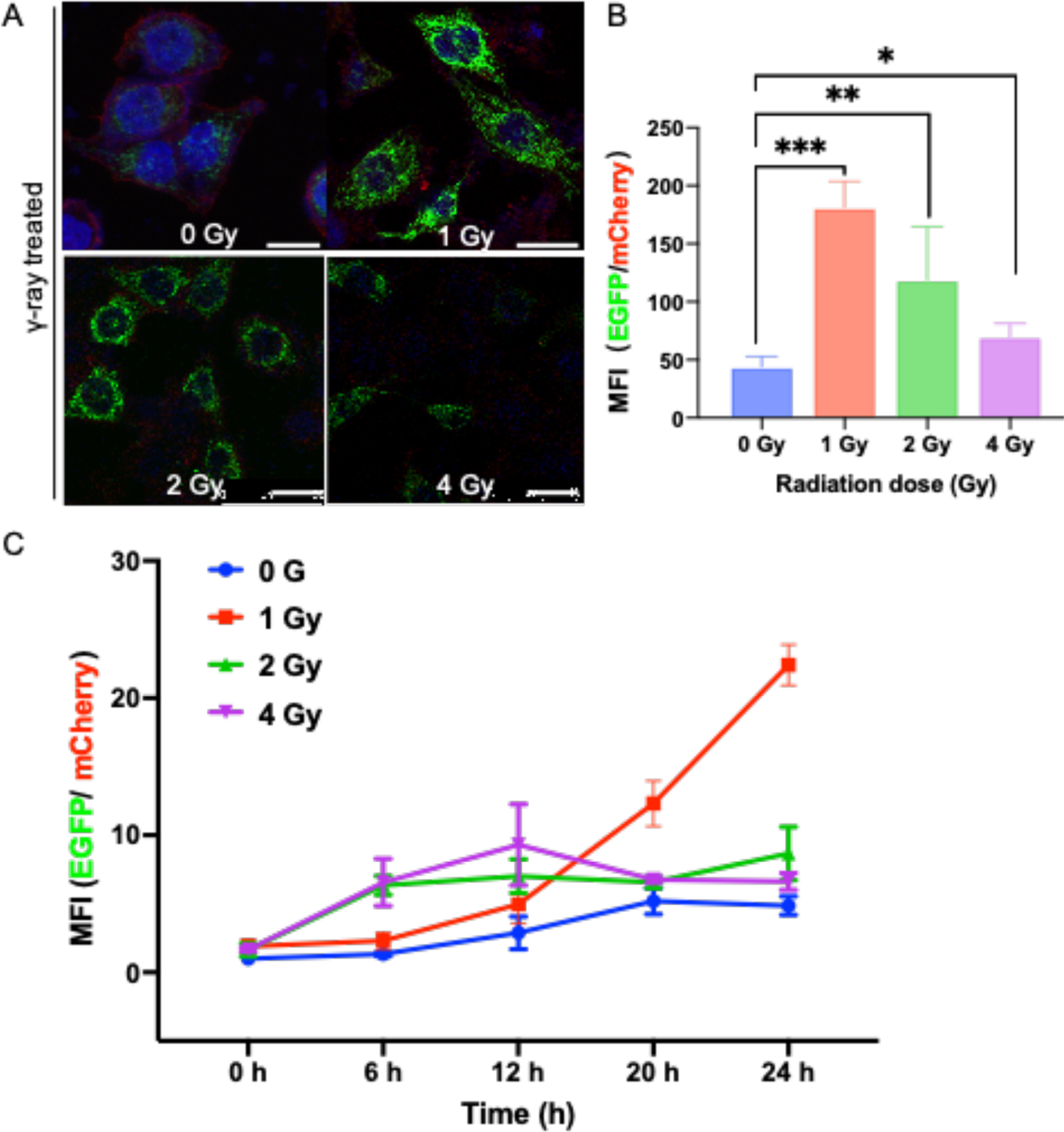
In-situ observation of the biological effects of gamma-radiation via the dual-fluorescence reporter system (A) HeLa-cox8-EGFP-actin-mCherry cells were exposed to 1 Gy, 2 Gy, 4 Gy gamma-ray and incubated for 24 h and observed with confocal laser scanning microscope. Green color indicates the mitochondria encoding COX8-EGFP, red color shows the actin encoded by actin-mCherry, and blue color represents the DAPI stained nuclei. Scale bar: 25 μm. (B) The ratio comparison for the mean fluorescence intensity (MFI) of GFP over mCherry fluorescence intensities quantified in the Figure 3A. The MFI was determined by ImageJ software. Bar graphs are presented as mean ± sd, n = 3. (C) The cellular mean fluorescence intensity (MFI) measured by Thermo High Content Imaging and Screening (HCS) system. HeLa-Cox8-EGFP-actin-mCherry cells were exposed to 1 Gy, 2 Gy, and 4 Gy gamma-ray respectively. MFI changed with time after the γ-ray irradiation.

Figure 3A presents the fluorescent images of the cells by confocal microscopy, indicating that the intracellular mitochondrial content increased with radiation dose. Compared with the control group (0 Gy), the irradiated group showed mitochondrial fluorescence enhancement, and it was significantly enhanced especially for the 1 and 2 Gy cases (Figure 3B). The fluorescent images intuitively show that the number of mitochondria increased with increasing radiation dose, confirming that the dual fluorescence reporter system can directly reflect the relevant mitochondrial responses and changes in the living cells.

To investigate the mitochondrial fluorescence enhancement of the cells exposed to ionizing radiation, the mean fluorescence intensity (MFI) from the dual fluorescence reporter system was utilized. This MFI ratio change was also obtained by employing the High Content Imaging and Screening (HCS) system, with the results shown in Figure 3C. The HCS living cell workstation recorded the changes of cell fluorescence within 24 hours after cell irradiation (the videos were also recorded, see Videos S1, S2, S3 in the Supporting Information). The results indicate that for 1 Gy, the fluorescence ratio GFP/mCherry increased with time, while for 2 Gy and 4 Gy cases, the fluorescence ratio increased significantly with the irradiation dose during the first interval of 1-12 hours, but afterwards, it decreased to some extent, though still higher than that of the control (non-irradiated cells). The reduction in MFI suggested that the mitochondria of the irradiated cells with higher radiation doses were actually more detrimentally damaged by the radiation than that received lower radiation doses. Figure S2 shows the numerical changes of two fluorescence channels detected by flow cytometry. The MFI ratio of EGFP^+^: mCherry^+^ was calculated, and it was found that the fluorescence ratio was positively correlated with the radiation dose. The result by the flow cytometry (Figure S2) also confirms there were significant differences between the irradiated and non-irradiated groups. To be noted, in order to verify the validity or reliability of actin fluorescence as an internal reference even under the irradiation condition, in-situ observation of real-time fluorescence data were also recorded after irradiation which showed that there was no significant difference in fluorescence expression within 24 hours (Figure S3).

### Radiation effect on cell viability, survival rate, cell cycle, apoptosis, DNA damage and mitochondrial membrane damage

To investigate the radiation effect, the cell viability was measured by CCK-8 method, showing that the cell viability decreased with the increase of radiation dose (Figure 4A). The toxic effect of radiation on cells was also examined using LDH kit, which revealed that the radiation-induced toxicity was indeed increased significantly with radiation dose (Figure 4B). Also, the colony-forming method was used to analyze the inhibitory effect of γ-ray on HeLa cell survival. With increase of γ-ray irradiation dose, the cell survival rate was only 30% after 4 Gy irradiation (Figure 4C), confirming that the cell survival rate decreased with the rise of radiation dose.

**Figure 4.**
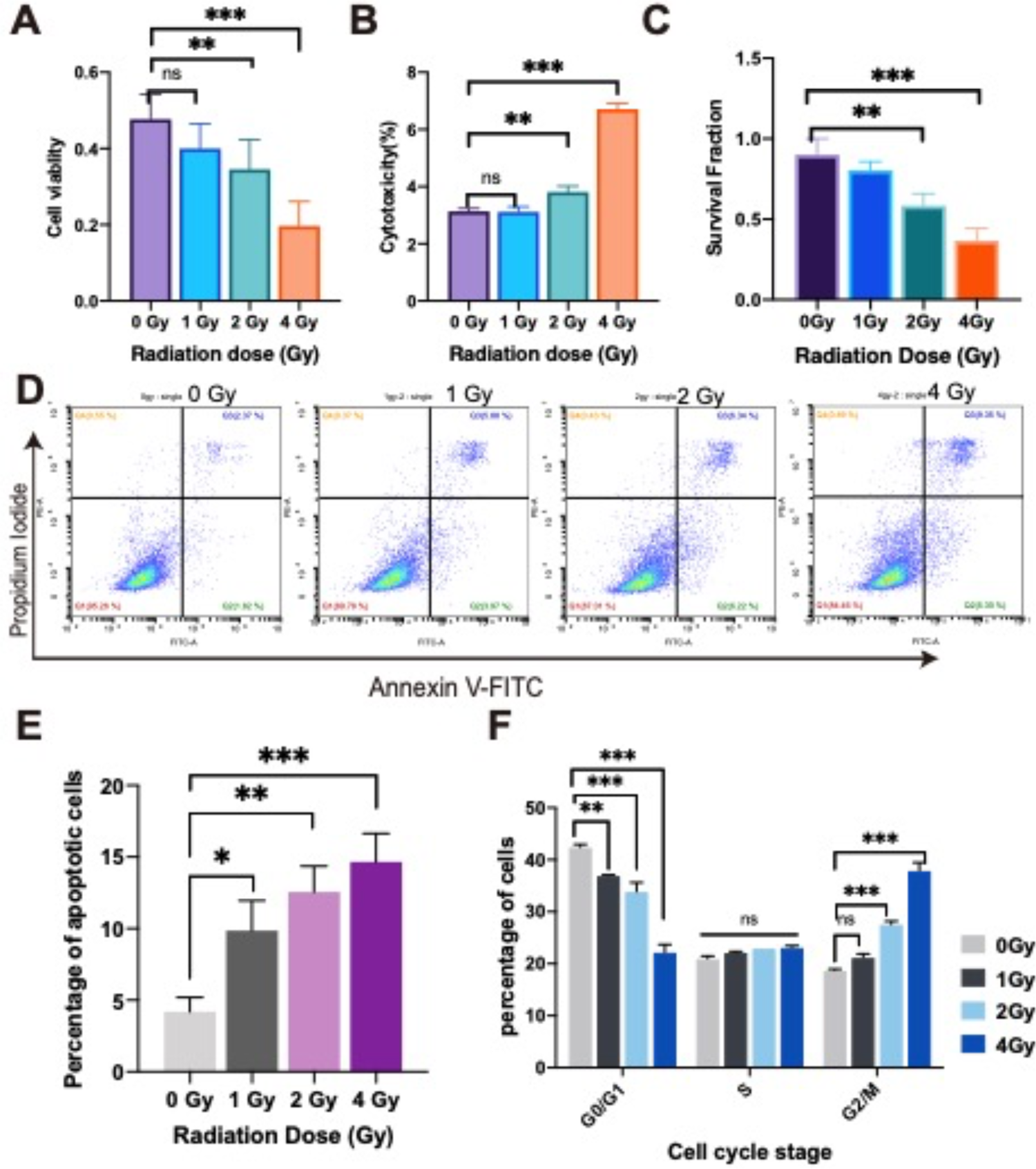
Detection of the radiation effect on HeLa cells exposed to different radiation doses (A) CCK-8 assay for cell viability. (B) LDH assay for cell cytotoxicity. (C) Colony formation assay of HeLa cells after 14 days under different radiation doses. (D) Flow cytometry results of Annexin V-PI assay. (E) Apoptotic cells at different radiation doses. (F) Cell cycle examination. The experiment was performed 3 times. Data are presented as Mean±SD; P < 0.01(**); P < 0.001(***); ns: not significant compared with untreated cells.

In addition, the cell cycle and apoptosis were assessed for the cells irradiated with γ-ray. Apoptotic cells were detected by flow cytometry after irradiation using Annexin V-PI methods (Figure 4D). Through statistics and comparison of the number of apoptotic cells, it was found that with the increase of radiation dose, the number of apoptotic cells also increased significantly (Figure 4E). Cell cycle was analyzed by flow cytometry, and the results showed that all groups of cells had G2 arrest after 24 hours of irradiation, especially at 2 Gy and 4 Gy irradiation doses showed significant G2 arrest (Figure 4F).

Furthermore, in order to explain the reasons for the radiation effects, the radiation induced damages on DNA and mitochondrial membrane, together with the accompanied cytosolic ROS (cROS) and mitochondrial-ROS (mROS) levels, were also examined, as shown by the results in Figure 5. In the experiments, the immunofluorescence method was used to detect DNA damage, CellROX Green probe was used to label cytosolic ROS, mitoSOX-Red probe was used for mitochondrial ROS level, and JC-1 was used to label mitochondrial membrane potential. Figure 5A shows that with increase of the radiation dose, the cytosolic ROS expression level increased significantly, and the statistical results showed that the ROS level in the irradiated cells increased with the increase of the radiation dose (Figure 5C). At the dose of 2 Gy, the cROS level was the highest, and for the 4 Gy group, there was a significant difference compared with the non-irradiated group. Besides, we also employed a mitochondrial biogenesis inhibitor, namely cyclosporin A (CsA), as for the negative control. Indeed, with the treatment of CsA, the cROS level was significantly increased (Figure 5C).

**Figure 5.**
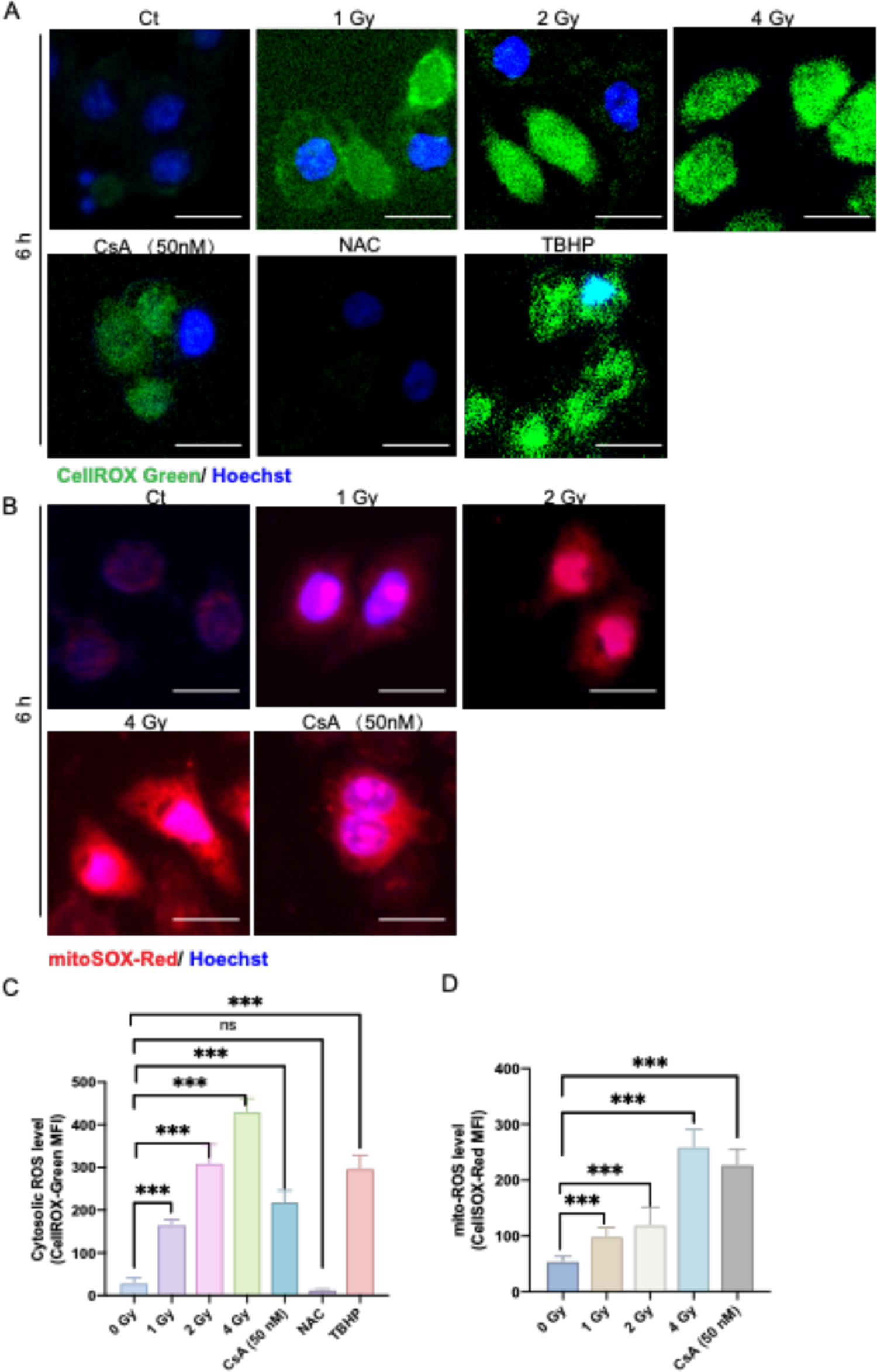
Analysis of cytosolic ROS (cROS) evel and mitochondrial ROS (mROS) level after irradiation of the cells (A) cytosolic ROS assessment using CellROX Green probe at 6 h. (B) mitochonrial ROS assessment using MitoSOX Red probe at 6 h. (C) The graph shows the representative MFI histogram of CellROX Green (cytosolic ROS production). (D) The graph shows the representative MFI histogram of MitoSOX (mitochondrial superoxide production). n = 500 cells calculated per group. Scale bar: 20 μm

ROS could induce mitochondrial damage, which would further activate the positive feedback circuit, resulting in more ROS production in damaged mitochondria. Therefore, we also used MitoSOX probe to detect mitochondrial ROS (Figure 5B). We observed an insignificant increase in the mitochondrial ROS level with increasing radiation dose compared with that of the control group (Figure 5D). Similarly, after treated with 50 nM CsA, the level of mitochondrial ROS was increased due to the inhibition of mitochondrial biogenesis. To be noted, mROS can be released into the cytoplasm through mitochondrial permeability transition pore (mPTP), which is also an important source of cROS in the cytoplasm. Therefore, we could expect that ionizing radiation would as a result also cause the increase of both mROS and cROS simultaneously.

For the mitochondrial damage evaluation, it is known that mitochondrial membrane potential (ΔΨ_m_) is produced by proton pump of electron transport chain, which is necessary for ATP production. Therefore, we also evaluated ΔΨ_m_ using JC-1 staining. As shown in Figures 6A-B, the promotion of JC-1 monomers enhanced significantly with the increase of radiation dosage. It was found that ΔΨ_m_ was markedly lower depending on the increased radiation dosage. After CsA (50 nM) treatment, the mitochondrial membrane potential decreased (Figure 6C).

**Figure 6.**
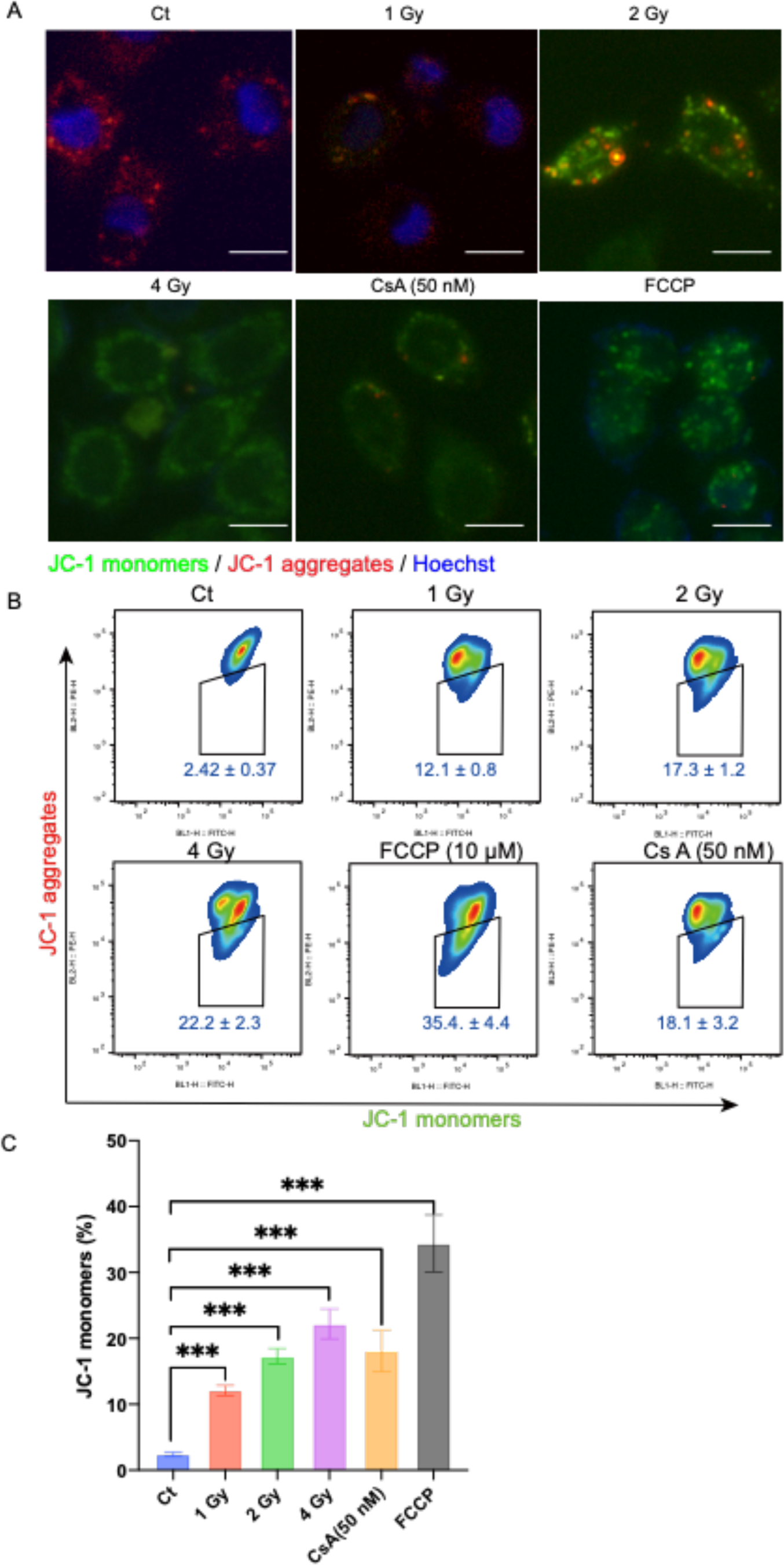
Analysis of mitochondrial membrane potential change after irradiation of the cells (**A-B**) Mitochondrial membrane potential change was assessed by JC-1 staining at 12h after irradiation. JC-1 monomers (green) and aggregates (red) were detected by fluorescence microscopy (**A**) and flow cytometry (**B**). scale bars: 10 μm. (**C**) The graph shows the proportion of JC-1 monomers change with different irradiation doses.

As the ionizing radiation induced DSB of DNA, we expected that there would also be enhanced DNA repair process in the irradiated cells. Indeed, we observed the fluorescence from γ-H2AX and the co-localization of 53BP1 foci after the radiation exposure. Figure 7 shows the number of foci increased significantly with the increase of radiation dose, while the number of foci of 53BP1 was the largest at 2 Gy but then decreased with radiation dose, implying that at higher radiation dose the cell received severer damage son the DNA repair ability became weaker.

**Figure 7.**
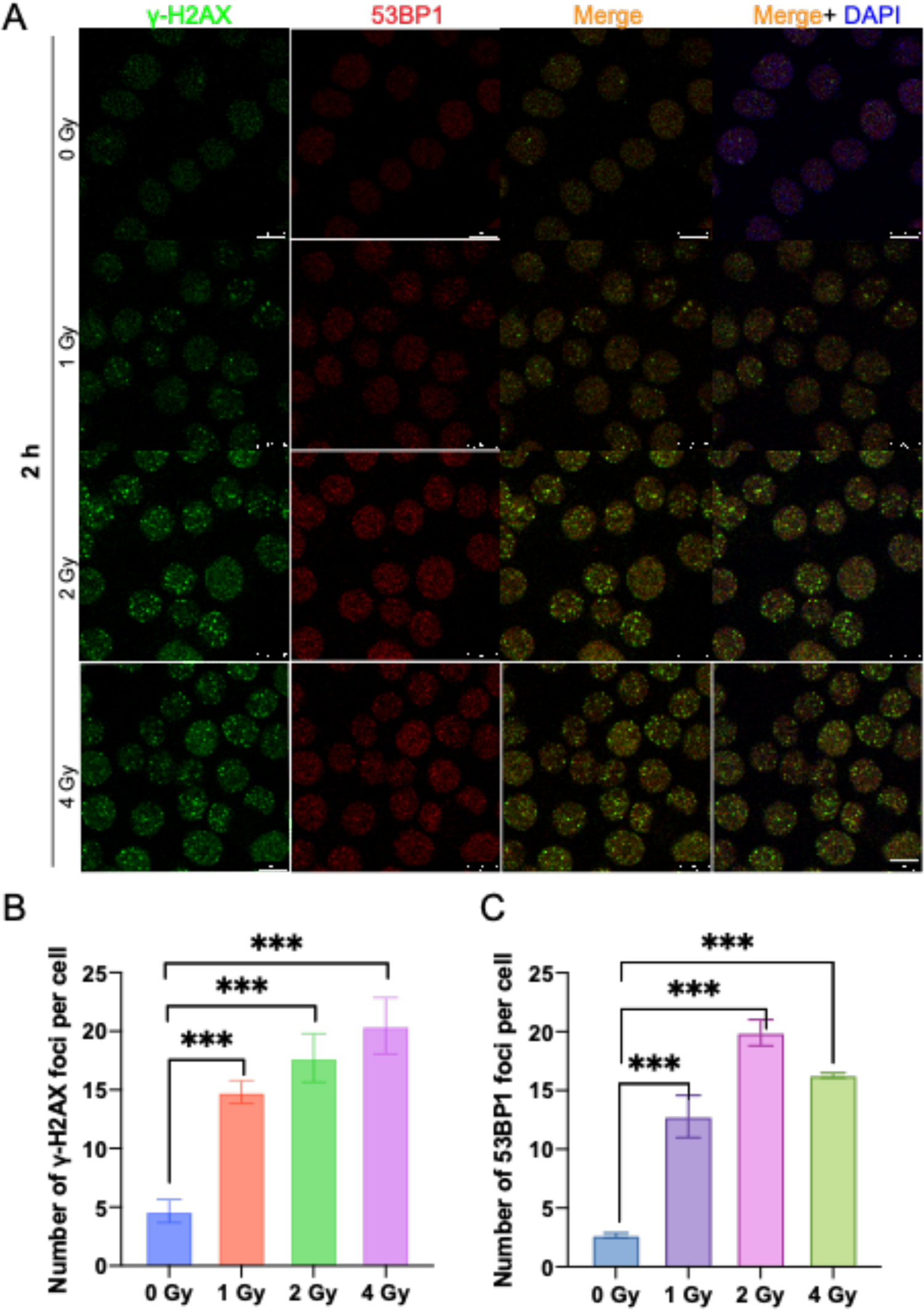
Comparative analysis of γ-H2AX and 53BP1 foci generated in the irradiated cells (A) Representative images for DNA damage detection when the cells were exposed to radiation with different radiation doses. Green: γ-H2AX; red: 53BP1; Blue: DNA nucleus stained with DAPI; Scale bars: 10 μm. (B) The percentage of population with the noted number of γ-H2AX foci corresponding to the respective fluorescence images. (C) The percentage of population with the noted number of 53BP1 foci corresponding to the respective fluorescence images. At least 100 nuclei per genotype were used for quantification. Image auto-quantification was performed with CellProfiler version 4 (www.cellprofiler.org).

### Verification of mitochondrial calcium and ATP level

Mitochondrial calcium plays a key role in regulating cell homeostasis, which has a dual regulation mechanism. On one hand, mitochondrial calcium can affect cell activity by activating oxidative metabolism, mitochondrial respiration and ATP synthesis. On the other hand, the increase of mitochondrial calcium influx is also one of the inducements of apoptosis and necrosis. In our experiment, we employed a cytoplasmic calcium probe (Fluo-4 AM) and a mitochondrial calcium probe (Rhod-2 AM) to detect the changes of calcium in the cytoplasm and mitochondria, and we obtained the result as shown in Figure 8A, which indicates that with the increase of radiation dose, the fluorescence intensity of Fluo-4 AM and Rhod-2 AM increased significantly. The analysis of the fluorescence intensities shows that the increase of calcium content in cytoplasm and mitochondria was positively correlated with radiation dose (Figures 8B-C). In addition, we treated the cells with CsA, which could inhibit the mitochondrial biogenesis, and the result confirmed that CsA also significantly improved the calcium release in both cytoplasm and mitochondria.

**Figure 8.**
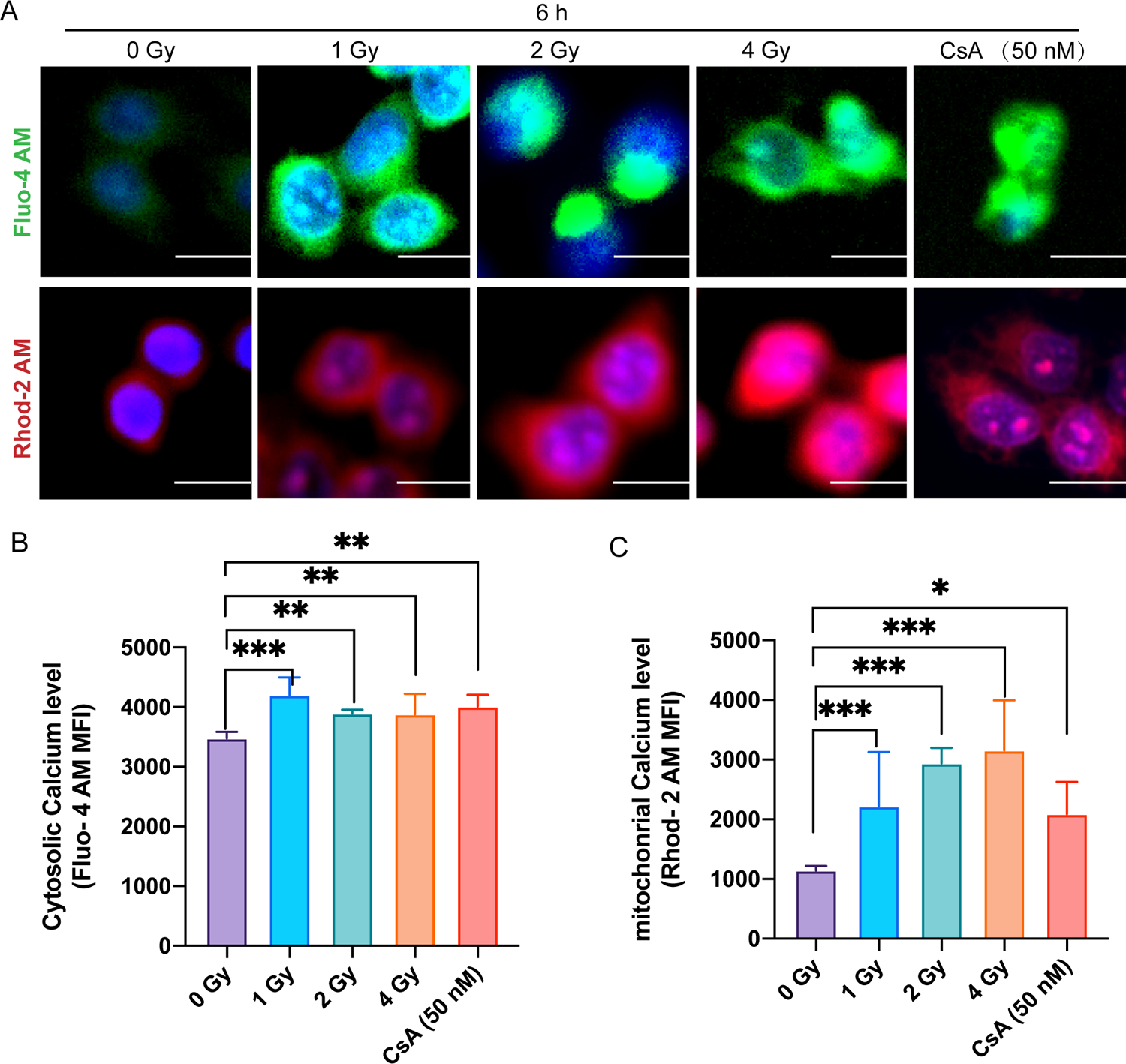
Radiation induces mitochondrial Ca^2+^ accumulation (**A**) Photomicrograph shows the radiation induced change and cellular distribution of Ca^2+^ stained with Fluo-4 AM and mitochondrial calcium using Rhod-2 AM. (**B-C**) Bar diagram shows alteration in Ca^2+^ concentration in HeLa cells stained with Fluo-4 AM (5 µM) and Rhod-2 AM (1 µM), respectively, analyzed using fluorescence microscope and imaging by imageJ software. Images were captured under fluorescence microscope with 20X objective. n = 500 cells calculated per group. Scale bar: 20 μm.

The imbalance of calcium homeostasis can further increase the production of reactive oxygen species and trigger the imbalance of mitochondrial energy metabolism (including the release of cytochrome c and apoptosis caused by permeability transition pore). The level of ATP is a marker to detect the level of mitochondrial energy metabolism, so we also detected the expression of ATP in cells at different time points (6, 12, 24 h) with different radiation doses (1 Gy, 2 Gy, 4 Gy). As shown in Figure 9, at each time point, with the increase of radiation dose, intracellular ATP level decreased significantly. For the same radiation dose, the expression level of ATP decreased significantly with the increase of time.

**Figure 9.**
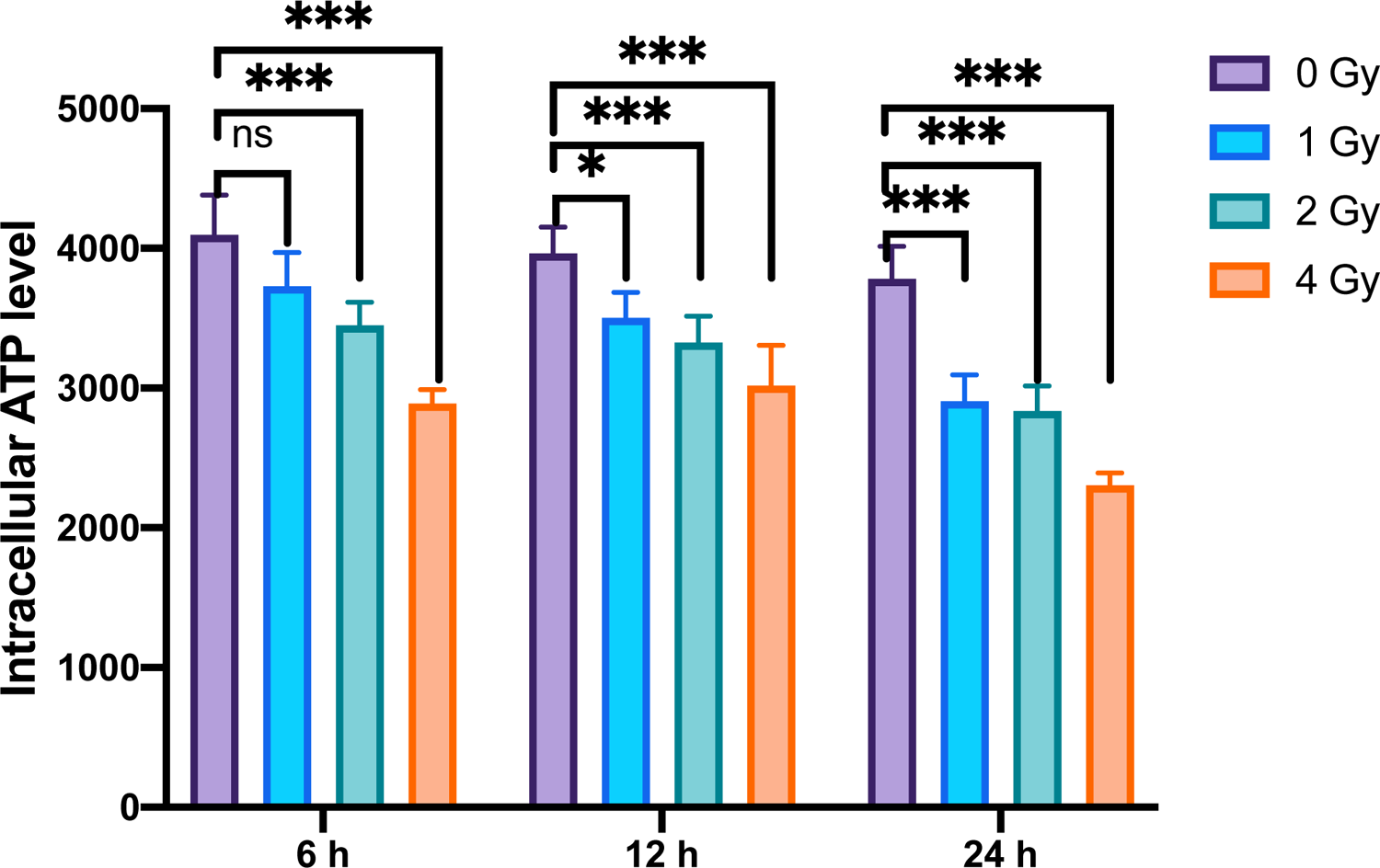
Effects of intracellular ATP depletion by radiation. HeLa cells treated with different doses of radiation (0, 1, 2 and 4Gy). Quantitative analysis of intercellular ATP level in percentage. *P<0.05, **P<0.01 and ***P < 0.001 indicated significant differences compared to non-treated cells group. All the results are presented as mean ± SD; n=3.

### Verification of mitochondrial biogenesis, apoptosis and mitophagy

To further verify the mitochondrial biogenesis induced by radiation, the copy number of mitochondrial DNA was evaluated firstly. Total DNA was extracted from HeLa cells 6-24 hours after irradiation. As shown in Figure 10A, the copy number of mitochondrial DNA changed significantly after cell irradiation. Indeed, under the same radiation dose, mitochondrial DNA number increased significantly compared with that of the non-irradiated cells. At 6 and 12 hours, the ratio of mitochondrial DNA to nuclear DNA increased significantly with the increase of irradiation dose. At 24 hours, the ratio was the highest at 1 Gy, and decreased with the increase of radiation dose.

**Figure 10.**
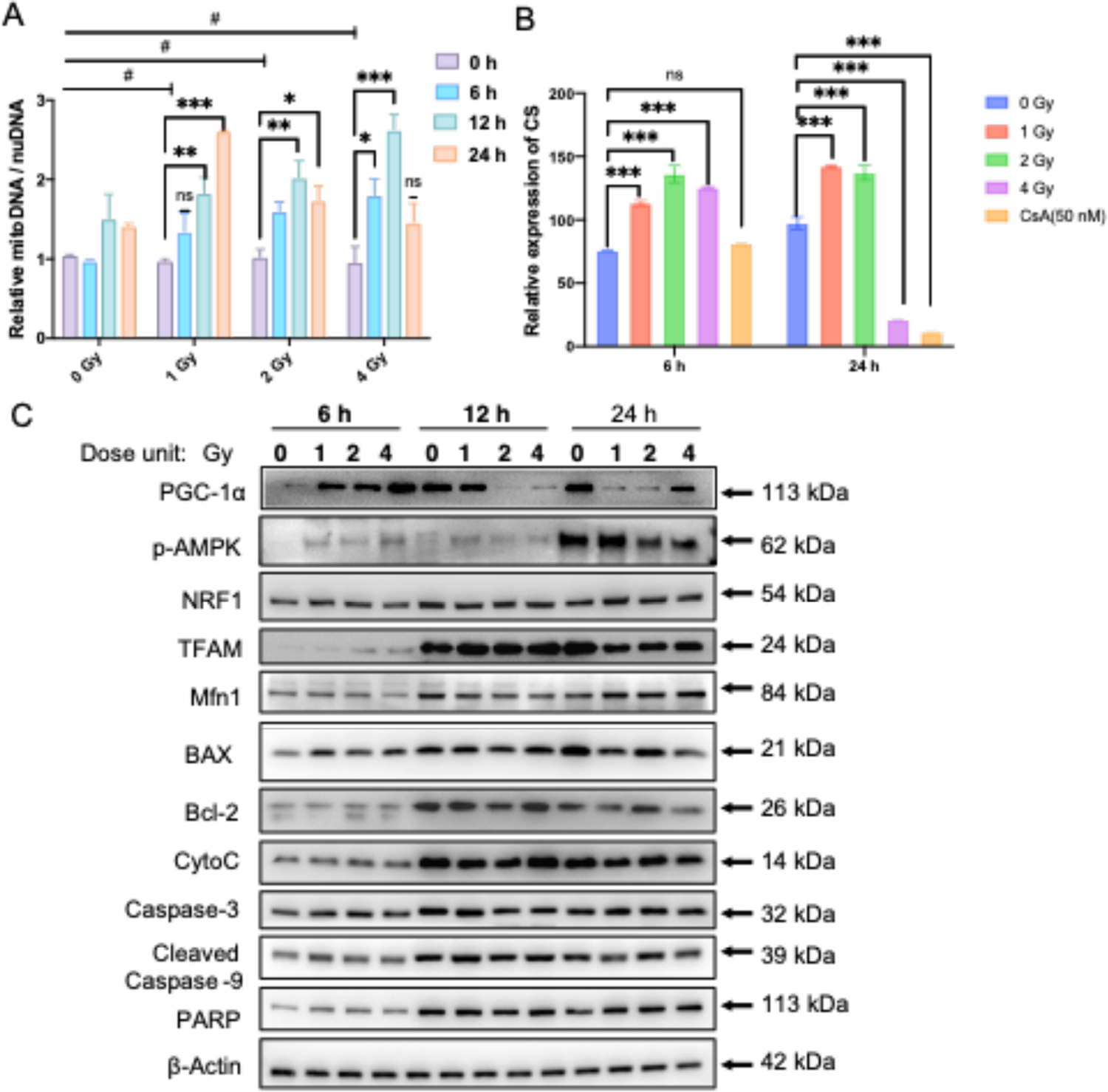
Radiation interferes with mitochondrial function and mediates apoptosis (A) Effect of gamma-gay irradiation on mitochondrial DNA copy number. HeLa cells were exposed to gamma-ray at doses of 1 Gy, 2 Gy and 4 Gy, respectively. After 6, 12, 24 hours, total genomic DNA was extracted and used as a template to detect the mtDNA/ nDNA ratio by RT-qPCR. Data are presented as Mean±SD; #: significant compared with unirradiated group; \*\**P* < 0.01; \*\*\**P*< 0.001. (B) Effect of irradiation on the activity of citrate synthase (CS). HeLa cells were exposed to gamma-ray at doses of 1 Gy, 2 Gy and 4 Gy, respectively. After 6, 24 hours, the expression level of citrate synthase activity was detected by UV-Vis spectrophotometer. (C) Western blot assay of the related proteins after radiation treatment. HeLa cells were treated with different doses of irradiation (0, 1, 2, 4 Gy), and total proteins were extracted at different time points. The WB assay shows the changes of expression of mitochondrial biogenesis marker proteins (PGC-1α, p-AMPK, NRF1, TFAM and Mfn1) and apoptotic marker proteins (BAX, Bcl-2, CytoC, Caspase-3, Cleaved Caspase-9, PARP) with time.

Citrate synthase (CS) is the initial enzyme of tricarboxylic acid (TCA) cycle. It exists in almost all cells that can be oxidized and metabolized, and it is also an important exclusive marker of mitochondrial matrix. Since CS enzyme is a key biomarker of mitochondrial biogenesis, we therefore also detected the activity of CS enzyme change after irradiation. As shown in the Figure 10B, at 6 hours, with the increase of radiation dose, CS increased significantly, suggesting that radiation stimulated mitochondrial biogenesis at the early stage. At 24 hours, low dose irradiation (1 Gy) could still increase the activity of CS, while it was reduced at higher radiation doses (2 Gy, 4 Gy), indicating that when the radiation damage became severer, mitochondrial biogenesis would diminish in the late stage of cell process. In addition, we also applied CsA as the inhibitor of mitochondrial biogenesis for the negative control. Indeed, after CsA treatment, the activity of CS decreased, and this decrease was more significant with increase of time.

Furthermore, to explore the correlation between mitochondrial biogenesis and apoptosis under the condition of radiation, the related marker proteins were also examined. For the mitochondrial change, the related protein expression levels of PGC-1α, NRF1, TFAM and Mfn1 at different times (6 h, 12 h, 24 h) with different radiation doses (0 Gy, 1 Gy, 2 Gy, 4 Gy) were assessed by Western blots (Figure 10C). At early stage (6 hours) of the process, the expression level of PGC-1α, p-AMPK, TFAM protein was increased with radiation dose. On the other hand, for the apoptosis, the related protein expression levels of apoptosis-related marker proteins (BAX, Bcl-2, CytoC, Caspase-3, Cleaved Caspase-9, PARP) were examined. As shown in Figure 8, the expression levels of crucial apoptosis proteins did not differ significantly at early time (6 h), but their expression levels increased significantly afterwards (12 h, 24 h). So these results unambiguously demonstrate that the mitochondrial biogenesis at early time was positively correlated to the apoptosis in the late stage.

So, as shown above, in the early stage after radiation, mitochondrial biosynthesis would be stimulated and increased, but with the lapse of time, for the larger irradiation dose, we found the decrease of mitochondrial energy metabolism (including the decrease of citrate synthase, mitochondrial membrane potential, mitochondrial DNA copy number, etc) and the inhibition of mitochondrial biogenesis. As the mitochondrial damage became severer, we also expected that mitophagy might play an important role to clear up the dysfunctional mitochondria, because mitophagy is an important degradation mechanism to eliminate dysfunctional mitochondria through autophagosomes(Montava-Garriga and Ganley, 2019). Therefore, we examined mitophagy by stably transfecting the mKeima plasmid to target the expression on the mitochondria of HeLa cells. mKeima fluorescent protein can change its excitation wavelength and show different color fluorescence at different pH values. Under pH neutral condition, it can be excited to green fluorescence at 485 nm, while under acidic condition, it can be excited to red fluorescence at 560 nm. We thus evaluated the intensity ratio of 560 nm to 485 nm fluorescence bands to determine the occurrence of mitophagy. As shown in Figure 11A, the ratio increased significantly with the increase of radiation dose, suggesting that at higher radiation dose mitophagy became dominant. AICAR, as a stimulant of mitochondrial biogenesis (positive control), also induced mitophagy, while CsA as an inhibitor of mitochondrial germinating (negative control), did not induce mitophagy (Figure 11B). To be noted, the mitophapy became most prominent at 9 hours after irradiation of the cells (Figure S4). These results indicate that the early-staged radiation-induced mitochondrial germination would be suppressed by the subsequent mitophagy, and if the mitochondrial damage was aggravated at higher dose radiation, the mitochondrial biogenesis was then diminished in the late-stage of the cell process.

**Figure 11.**
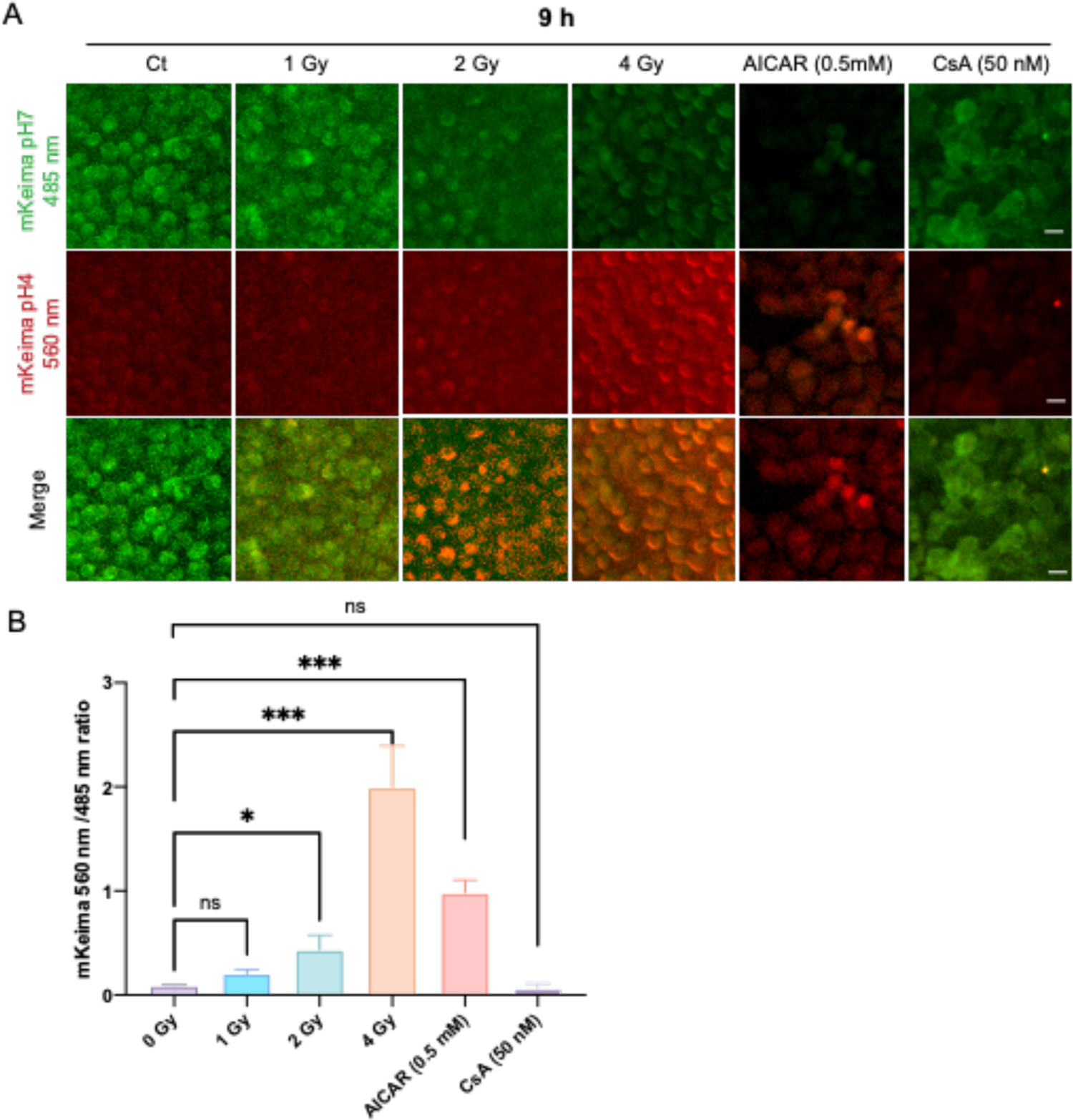
Mitophagy activity assessed using mKeima in HeLa cells Elevated levels of mitophagy were observed following irradiation with 1 Gy, 2 Gy, 4 Gy after 9 hours. AICAR (0.5mM) and mitophagy inhibitor (CsA, 50 nM) were also applied as the positive and negative controls. n = 500 cells calculated per group. Scale bar: 20 μm.

## DISCUSSION

Mitochondria as the cell energy factory play a vital role in cell homeostasis(Lill and Freibert, 2020). In response to external stimuli, mitochondrial biogenesis can be elicited in order to provide the cells with extra energy(Barros and McStay, 2019). Previously, many studies had reported that mitochondrial biogenesis could suppress apoptosis(Yu et al., 2013), i.e., apoptosis is negatively correlated with mitochondrial biogenesis. Some studies even claimed that apoptosis occurred because of the absence of mitochondrial biogenesis(Martins et al., 2015). But this understanding is not accurate, because to the best of our knowledge, mitochondrial biogenesis should be occurring as the early event prior to apoptosis, though it might be reduced or diminished later during the process of cell apoptosis. So, to explore the relationship between mitochondrial biogenesis and apoptosis, it is important to scrutinize the whole cell process and examine the respective bio-effects at different time points. Would mitochondrial biogenesis occur preceding the apoptosis? And, if there would be mitochondrial biogenesis occurring as the early event of the apoptosis, how would it change with time? With these questions and considerations, we thus initiated this research, and applied ionizing radiation as the tool to induce both mitochondrial biogenesis and apoptosis in the cells, to explore the time-dependent relationship between mitochondrial biogenesis and apoptosis. As a result, we have thus achieved the following results with some new findings and elaborated understandings.

Firstly, we have constructed the dual fluorescence report system that can be effectively used for mitochondrial biogenesis study. To probe the changes taking place in mitochondria, we have especially developed the approach using mitochondrial-tagged recombinant fluorescent proteins. To the best of our knowledge, it is actually the first time to apply the dual fluorescence ratio approach to study the biological effects of mitochondria. According to the in situ and real-time observation of fluorescence in the cell workstation and the high-throughput analysis of fluorescence ratio through high-content quantitative analysis system, we have thus developed a new method of using dual fluorescent reporter system for the the study of mitochondrial biogenesis, which has been proved to be quite effective and reliable. As we know, the fluorescent protein reporting system has the advantages of real-time, convenient, fast, and non-destructive(Betzig et al., 2006). Besides, it has also an advantage in the study of subcellular organ localization and protein stress regulation(Llopis et al., 1998). Previously, people often used fluorescent dye probes for studying effect involved with mitochondria. For example, Mukhopadhyay et al. used Annexin V and Sytox Green and MitoSOX Red fluorescent probes to study the detection of apoptosis and mitochondrial superoxide so as to establish the method for detecting living cells by laser confocal scanning microscopy and flow cytometry (Mukhopadhyay et al., 2007). But the disadvantages of fluorescent dyes are the toxicity to cells, as well as the light quenching under long-period real-time observation. Compared to fluorescent dyes, fluorescent proteins are more stable and have minimal toxicity and can generate visible fluorescence in vivo (Westermann and Neupert, 2000). Actually, GFP reporter protein is now commonly used to detect mitochondrial changes(Hanson et al., 2004; Mahajan et al., 1998). In this work, in particular, we have constructed the dual fluorescence reporting system based on fluorescence proteins for monitoring the mitochondrial biogenesis process. We have also verified the effectiveness of this system designed for monitoring mitochondrial biogenesis. For this purpose, we apply AICAR to activate mitochondrial biogenesis as the positive control (Komen and Thorburn, 2014), as AICAR is an activator of AMP-activated protein kinase (AMPK) that can permeate cell membranes(Rai et al., 2015). AMPK (AMP activated protein kinase) is an activator of mitochondrial biogenesis and key regulator of energy homeostasis caused by external factors (Kim et al., 2016). The main function is to phosphorylate PGC-1α protein, or affect the transduction of SIRT1 signaling pathway(Mihaylova and Shaw, 2011). Our result clearly shows that AICAR indeed induced the up-regulation of key protein levels of mitochondrial biogenesis. The prominent merit of such a dual fluorescent protein reporter system is that it is not only suitable for the real-time in situ observation for a long time, but also facilitates the precise fluorescence evaluation with the reference fluorescent protein for intensity normalization, so that the interference from background signal noise can be best avoided. Therefore, by analyzing the ratio of the fluorescence intensity of mitochondrial DNA to that of nuclear DNA expression, the early event of mitochondrial biogenesis under irradiation condition can be readily observed and evaluated, and with this approach we can thus sensitively track the early process that which would eventually lead to the late event of apoptosis.

Secondly, with the as-constructed dual fluorescence report system, we have observed the radiation-induced mitochondrial biogenesis together with the apoptosis. Ionizing radiation has been used for cancer therapy for they can directly or indirectly cause DNA damage and kill the cancer cells(Spitz et al., 2004). With DNA damage in the nucleus, cells may undergo cell cycle arrest required for repairing the damage or cell death including apoptosis(Bernstein et al., 2002). In parallel, mitochondria is also considered to be involved in the radiation induced effects, where more reactive oxygen species (ROS) would be produced which may affect the cells profoundly. Actually, it has been well documented that exposure of cells to ionizing radiation could activate ROS to produce oxidases, regulate antioxidants, and alter metabolic activity in response to oxidative damage(Azzam et al., 2012). In both the nucleic or mitochondrial processes, more ATP is required to elicit mitochondria biogenesis either directly or indirectly(Kulms et al., 2002; Srinivas et al., 2019). So, it is not surprising that mitochondria biogenesis could take place as one of the consequences of radiation-induced bio-effects (Galluzzi and Kroemer, 2008). But interestingly, in our fluorescence measurements, we found that the fluorescence ratio increased and reached the maximum at 6 h after irradiation, but afterwards the fluorescence ratio decreased gradually. According to our established mitochondrial fluorescence report system, this suggests that in the early stage, mitochondrial biogenesis was indeed initiated and enhanced. In this process of mitochondrial biogenesis, mitochondrial DNA and mitochondrial mass were maintained to keep the cell homeostasis and metabolism(Huangyang et al., 2020). But in the late stage of the radiation effect, the fluorescence ratio decreased, showing that the function of mitochondrial biogenesis gradually diminished. We can also understand this change by relating it to the ROS elevation. There was increase of ROS caused by radiation, which continuously increased the oxidative stress and damaged the mtDNA and the mitochondrial integrity. For the role of ROS, it is generally accepted that normal range of ROS may induce stress response by changing the expression of related nuclear genes, and certain level of ROS can save cells by maintaining energy metabolism (Srinivas et al., 2019); but when the ROS level exceeds the tolerant threshold, ROS can then induce the transition of mitochondrial membrane permeability, leading to the activation of the caspase pathway and promoting cell apoptosis. In our experiment, we detected the increase of ROS level in cytoplasm and mitochondria, the increase of calcium ion level in cytoplasm and mitochondria, the decrease of ATP level and mitochondrial membrane potential, the decrease of mitochondrial DNA copy number and mitochondrial citrate synthase activity. All these results are consistent with each other. Also, given the increase of radiation dose, we detected the related caspase-related proteins and found that the increase of pro-apoptotic proteins such as cytochrome c. Release of cytochrome c is an indicator for the apoptosis, in which the oxidative stress would damage mtDNA and other components of the cells(Lee and Wei, 2005). Here, we especially notified that the mitochondrial fluorescence intensity change rates were quite different after the cell exposure to the radiation. The larger the irradiation dose, the higher the fluorescence ratio reduction rate. All these observations are actually the typical features for the radiation-induced apoptotic process.

Thirdly, our new observation has revealed the more subtle but very important relationship which had largely been substantially or largely ignored by previous studies. We noticed that in the beginning of the process, the mitochondrial biogenesis was actually positively rather negatively associated with the apoptotic stress, and the trend of fluorescence change was consistent with the number of apoptotic cells. This is actually contrary to the conventional view on the relationship between mitochondrial biogenesis and apoptosis. In the past, there were few reports concerning the dynamic process of mitochondrial biogenesis, and the mainstream opinion or understanding on the relationship between mitochondrial biogenesis and apoptosis was that these two processes normally appear to be antagonistic. For example, many studies showed that the apoptosis occurred simultaneously with the suppression, or, that apoptosis took place with even missing the observation of mitochondrial biogenesis (Lin et al., 2020; Martins et al., 2015; Vayssiere et al., 1994; Zhang et al., 2017). This is actually not true or accurate, because mitochondrial biogenesis as an earlier event did occur preceding the apoptosis, as we have clearly demonstrated in this study.

The reason for this misleading of understanding is largely due to the ignorance of the investigation of the dynamic process of the cells. Generally, people tend to choose just several time check points for inspecting the apoptosis and mitochondrial biogenesis. But which time check points are most suitable for the correct evaluation? What cautions should be taken to avoid the wrong evaluation when especially concerning the relationship between the processes in the involvement? To solve this, we actually need to establish more precise method to study the cellular effect in its dynamic process, and so to scrutinize the processes of both mitochondrial biogenesis and apoptosis in their logical time sequence. This is also one of the main purposes of this study. It is understandable why people had not focused on the study of processes, as we realized that there had been lack of tool/method for the monitoring the whole process regarding the mitochondrial changes in situ and in a nondestructive way. For example, the traditional methods include using real-time quantitative PCR to detect the copy number of mitochondrial DNA, or using Western blot assay to identify the changes of marker protein level of mitochondrial biogenesis. These methods cannot achieve in-situ and real-time observation in living cells. On the contrary, the fluorescence protein method can be used to observe the mitochondrial biogenesis in real time as especially useful for studying the early events prior to apoptosis. Therefore, in this work, we for the first time attempted to construct the fluorescent reporting system which could let us to track the change of mitochondrial biogenesis. Based on the fluorescence signal, the dual fluorescence ratio can help us to predict the radiation effects or the cell fate according to the fluorescence change of these irradiated cells.

In particular, from the fluorescence data obtained from Figure 3C, we can calculate the fluorescence ratios, and so we can retrieve the MFI ratios in each irradiation dose group (0 Gy, 1 Gy, 2 Gy, 4 Gy) at an early time, as shown in Figures 12 A-B. Strikingly, this result is consistent with the cells in the apoptotic state at 24 h (Figure 4E). In the late stage (24 hours), the fluorescence ratio becomes smaller as the irradiation dose increases. It is understandable because the apoptosis is actually dependent on the early biochemical events in the cells(Holler et al., 2000). In this way, we may therefore infer or predict the late apoptotic state of cells by observing the early-staged changes in terms of mitochondrial fluorescence ratio. On the other hand, it is also seen that this fluorescence ratio was significantly reduced after 24 h (Figure 12C), which is not concomitant with the apoptosis at 24 h (Figure 4E). This suggests again that it is very critical to evaluate both the processes by choosing the right time check points.

**Figure 12.**
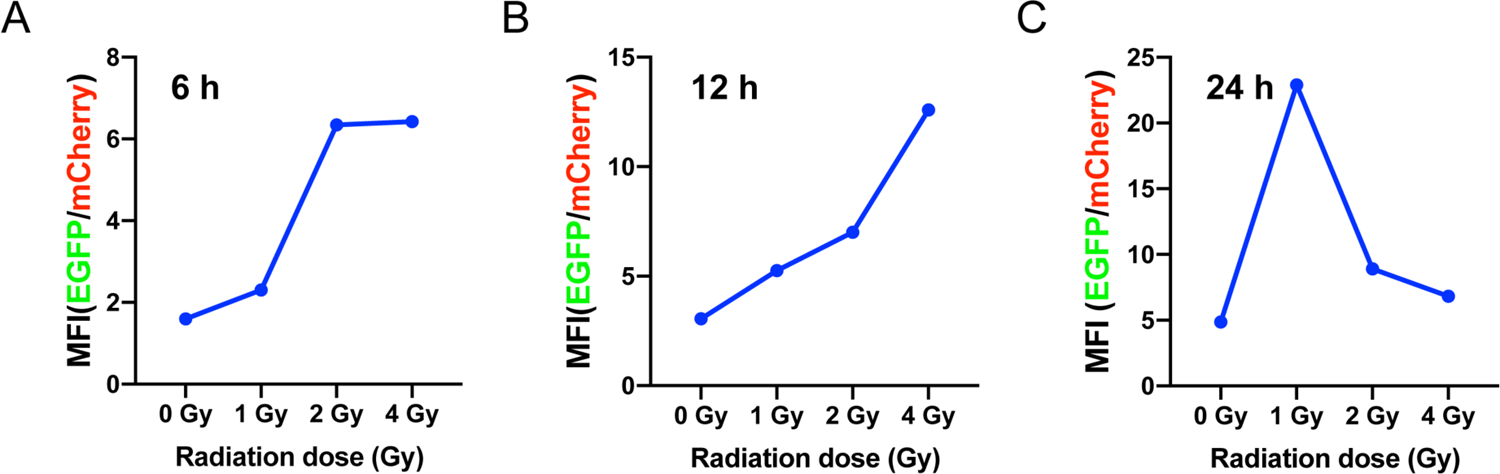
Analysis of mitochondrial biogenesis and apoptosis (A) Analysis of EGFP/mCherry ratio in High Content Screening (HCS) at 6 h. (B) Analysis of EGFP/mCherry ratio in High Content Screening (HCS) at 12 h. (C) Analysis of EGFP/mCherry ratio in High Content Screening (HCS) at 24 h.

Finally, we have also tried to understand the mechanism for the mitochondrial biogenesis related to the radiation-induced apoptosis. Generally, it is believed that the biogenesis of mitochondria is a very complex biological process, which can maintain the cell homeostasis by controlling the self-renewal of subcellular organelles and maintaining mitochondrial DNA (Scarpulla, 2008). The initiation of mitochondrial biogenesis is normally triggered by the reception of energy demand signals caused by external stress (Popov, 2020), while at the same time, mitochondrial biogenesis itself is a protection way for cell’s self-renewal. Since it is well-known that ionizing radiation is an effective way to stimulate the cell mitochondrial biogenesis, we thus employed the tool of ionizing radiation to treat the cells. As a result, we have successfully proved that some marker proteins of mitochondrial biogenesis are involved in the biological effects of radiation. We have found that the protein expression levels of nuclear respiratory factor1 (NRF1), peroxisome proliferative activated receptor, gamma, coactivator 1 alpha (PGC-1α), transcription factor A, mitochondrial (TFAM) increased with radiation dose. NRF1 is a transcription factor with a vital role in mitochondrial functional genes(Virbasius et al., 1993). As a transcription coactivator, PGC-1α is a central regulator of mitochondrial biogenesis and energy metabolism, and has become a hot target for cancer treatment because of its relationship with cell death and oxidative metabolism(Wang et al., 2019) (Bost and Kaminski, 2019) (Wu et al., 1999). TFAM is a key activator of mitochondrial (mt) DNA transcription. mtDNA is reported to be very susceptible to oxidative stress(May-Panloup et al., 2005). With the confirmation of these factors, we have thus also again unambiguously confirmed that this mitochondrial biogenesis is indeed related to radiation-induced cell adaptation and apoptosis.

But how to explain the suppression or disappearance of mitochondrial biogenesis for the higher dose of radiation observed in the late stage of the cell process which corresponds to the apoptosis? In our experiments, we observed that mitochondrial biogenesis increased in the early stage of the cell process, but with the increase of radiation dose, the mitochondrial biogenesis decreased substantially in the late stage. Firstly, we speculate that this change of mitochondrial biogenesis is related to DNA repair ability, and with increased radiation damage, the DNA repair mechanism was destroyed. It was previously reported that mitochondrial biogenesis can enhance DNA repair ability, and mitochondrial biogenesis is closely related to DNA repair ability(Li et al., 2020) (Fu et al., 2008). For example, it was reported that doxorubicin, a drug which can induce DSBs of DNA as accompanied by mitochondrial biogenesis (Kluza et al., 2004). Normally, there are two main repair pathways of DSBs in mammalian cells: non homologous end joining (NHEJ) and homologous recombination mediated repair (HR) (Löbrich and Jeggo, 2005). Mitochondrial biogenesis can promote DNA repair by inducing chromatin changes in DSB and regulating the activity of DNA repair factors, and this upstream factor has been found to be the silent information regulator (SIRT) (Paredes and Chua, 2016). SIRT is a family of sirtuins that depend on NAD^+^ lysine deacetylases, which may play an important role in mitochondrial biogenesis and cell metabolism(Chalkiadaki and Guarente, 2015). On the other hand, DSB repair can be achieved by NHEJ which can activate DNA-PK (Davis et al., 2014), while DNA-PK can activate AMPK signal pathway, which is just closely related to mitochondrial biogenesis (Park et al., 2017). In our experiment, we have indeed observed the change of DNA repair as reflected by the change of 53BP1, and this change is just concomitant with the mitochondrial biogenesis. Secondly, we also speculate that the mitochondrial autophagy induced by radiation occurred, which subsequently cleared up the ROS-damaged and dysfunctional mitochondria. According to the analysis of the mitophagy result, we found that mitophagy increased with radiation dose at 9 hours after the cell irradiation, although 24 hours after the irradiation, this process also diminished with the death of the cells.

So, with the above findings and discussion of mechanisms, we can therefore concisely describe the process of radiation affecting cells as schematically depicted in Figure 13. After a certain dose of radiation, DNA damage occurs, and the cell cycle is blocked in order to initiate the repair process. At the same time, radiation produces a large amount of ROS in the cell and affects the homeostasis of the cell. To resist the radiation stress or damage, mitochondria then start to provide more energy for DNA repair and maintenance of cell metabolism and homeostasis, leading to mitochondrial biogenesis. Next, with the continuous increase of ROS in cytoplasm and mitochondria, the increase of calcium ions released to mediate the imbalance of mitochondrial energy metabolism homeostasis, the decrease of ATP level and the decrease of citrate synthase activity, the decrease of mitochondrial membrane potential which eventually led to the release of Pro apoptotic protein cytochrome c and the activation of apoptotic caspase pathway, the programmed cell death was elicited, with the diminishing of mitochondrial biogenesis in the late stage of the cell process.

**Figure 13.**
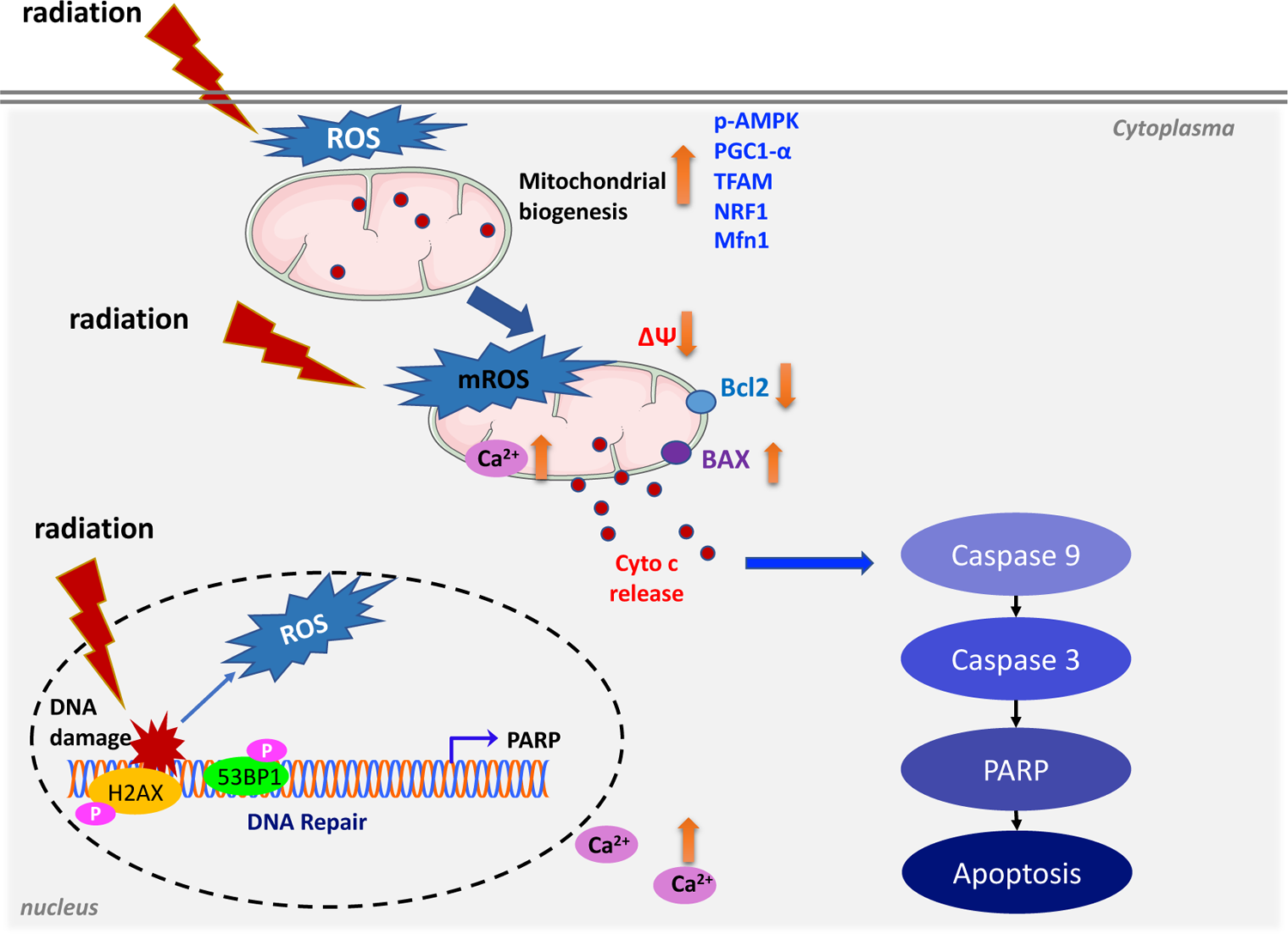
Schematic diagram of biochemical events after ionizing radiation

## CONCLUSION

In this work, we have constructed a dual fluorescence reporting system, which makes use of the intensity ratio of multiple fluorescence tags to observe the changes of mitochondria in real time, and these changes are related to ionizing radiation-induced mitochondrial biogenesis and apoptosis. We have also demonstrated that ionizing radiation can effectively cause mitochondrial biogenesis, so that it can be an efficient tool for the study of mitochondrial biogenesis. Particularly, for the exploration of the involved mechanisms, we have scrutinized the relationship between this mitochondrial biogenesis and apoptosis by considering the time sequence of occurrence. Our results have unambiguously proved that these two processes are not antagonistic if they are evaluated at right time points, but on the contrary, they may be positively correlated if we relate the mitochondrial biogenesis taking place at the early time to the apoptosis occurring at the late stage. Therefore, this work not only provides an effective method for directly observing the mitochondrial biogenesis process in living cells, but also demonstrates how to evaluate the biological effects correctly with right timing, with which we may be able to predict the cell fate such as apoptosis through the observation of the early event of cell process.

## STAR METHODS

### RESOURCE AVAILABILITY

#### Lead Contact

Further information and requests for reagents may be directed to the lead contact, Qing Huang.

#### Materials Availability

Reagents or materials used in this work may be requested from the Lead Contact by signing a completed material transfer agreement.

#### Data and code availability

Original data generated by this study are available upon request.

### EXPERIMENTAL MODEL AND SUBJECT DETAILS

#### Reagents

The *Escherichia coli* DH5a and DNA Ladder were purchased from Transgen (Beijing, China). *Nhe*I, *Hind*III, *EcoR*I, *Not*I restriction enzymes, T4 DNA Ligase, and Q5 High-Fidelity DNA Polymerase were purchased from NEB (MA, USA). Plasmid pEGFP-N1 was purchased from Clonetech Biotechnology (CA, USA). Plasmid pLVX-mCherry, PUC57 were stored in our laboratory. Plasmids of psPAX2 (# 12260) and pMD.2G (#12259) were purchased from Addgene. PCR purification and gel extraction kits were obtained from Axygen (CA, USA). TRIzol™ reagent, Lipofectamine 2000 were purchased from Invitrogen (CA, USA). Primescript II 1st strand cDNA Synthesis Kit, RT-qPCR kit and Total genomic DNA extraction kit were purchased from TAKARA (Dalian, China). G418 and Hoechst 33342 were purchased from Targetmol (Shanghai, China). Citrate Synthase Assay Kit was purchased from Solarbio (Beijing, China). Reactive oxygen species (ROS) probe (CellROX Green and MitoSOX Red) was purchased from Invitrogen. JC-1 dye was purchased from Beyotime. 5-Aminoimidazole-4-carboxamide ribotide (AICAR) was purchase from Selleck (MA, USA). The antibodies of PGC-1α, NRF1, TFAM, Mfn1, BAX, Bcl-2, CytoC, Caspase-3, Cleaved Caspase-9, PARP, γH2AX(S139), GFP, p-AMPK were purchased from Bimake (MA, USA), 53BP1, anti-β-Actin antibody was purchased from CST (MA, USA). Alexa Fluor® 488 Goat Anti-Mouse (IgG) secondary antibody, Alexa Fluor® 647 Goat Anti-Rat (IgG) secondary antibody were purchased from Invitrogen (CA, USA).

## METHOD DETAILS

### Creation of stably transfected cell lines

#### Generation of plasmids

Firstly, the pEGFP-N1-COX8 expression plasmid was constructed. Briefly, the DNA sequence of cytochrome c oxidase 8A subunit was found from Genbank (NM_004074.2) (Hendrickson et al., 2010). Then, two complementary oligonucleotide strands were obtained by amplification using PCR, and the restriction sites of the *Nhe*I and *Hind*III were added at both ends. The sequences of oligonucleotide strands were listed in Table S1 (Supporting information). The DNA sequence (COX8A) was ligated on the PUC57 plasmid. Then, the appropriate oligonucleotides were annealed and ligated into pEGFP-N1 cloning cut with *Nhe*I and *Hind*III. After incubation overnight at 37°C, the target plasmids were extracted from the bacterial fluid using the EndoFree Maxi Plasmid kit.

Secondly, the PLVX-mCherry-Actin expression plasmid was constructed. Total RNA was extracted from HeLa cells by Trizol reagent and served as the template for cDNA synthesis, and then it was reversely transcripted using Primescript II 1st strand cDNA synthesis kit. The primers were designed according to the sequence of actin (Genbank no. NM_10277): *EcoR*I tailed forward (5′-gtcGAATTCatggatgatgatatcgccgcgc-3′) and *Not*I tailed reverse (5′-TATGCGGCCGCctagaagcatttgcggtggac-3′) primers which are named as Fw-actin-EcoRI and Rv-actin-NotI, respectively. The length of the amplified product was 1128 bp. The PCR product was separated by electrophoresis, and the target fragment was recovered after gel cutting and purification. The target gene fragment (actin) and the expression vector (pLVX-mCherry) with the same restriction site (*EcoR*I and *Not*I) were digested by restriction enzymes respectively, and then ligated with T4 ligase and transformed into *E. coli* DH5a for plasmid amplification.

#### Production of lentiviral supernatant and transduction of COX8-EGFP-mCherry-ACTIN-HeLa

Briefly, positive monoclonal cells (COX8-EGFP-HeLa cells) were constructed. When HeLa cells grew to the concentration of 80% in complete medium, they were then transfected with pEGFP-COX8 plasmid with lipofectamine 2000. After 24-48 h of transfection, G418 was added to screen positive monoclonal cells.

The lentiviral supernatant was prepared according to the protocol provided by addgene (http://www.addgene.org/tools/protocols/plko/#A). When the confluence of HEK293T cells cells in the 6-well plate reached 80%, Lipofectamine 2000 was used to transfect the pLVX-mCherry-ACTIN plasmid and the viral packaging helper plasmid psPAX2, pMD.2G into the cells. After incubation for 8h, the medium was replaced with the fresh medium containing serum, and the supernatant was collected 24 hours and 48 hours later. The virus supernatant was filtered through a 0.45 µm pore so as to infect COX8-EGFP-HeLa cells. Next, the COX8-EGFP-HeLa cells were cultured in a 6-well plate to 70% confluence, then the medium was removed, and the virus supernatant was mixed with fresh medium containing 8 μg/ml polybrene in equal proportions and added to the culture plate. The fresh medium was replaced after 24 hours and the medium containing 1 μg/ml puromycin was changed at 48 hours to start screening to obtain stable cell lines.

#### Cell Culture, Synchronization and Gamma irradiation

HeLa, HEK-293T cell lines were purchased from American Type Cultures Collection (ATCC, Manassas, VA). The HeLa and HEK-293T cells were cultured in Dulbecco’s modified Eagle’s medium (DMEM, HyClone) containing 10% fetal bovine serum (FBS, Gibco BRL). All cells were maintained at 37 °C in 5% CO2.

The cell cycle synchronization method was detailed in the previous work (Rodríguez-Martínez et al., 2020). The cells were incubated with thymidine (2.5 mM) for 18 hours, washed once with PBS buffer and replaced with fresh medium for 14 hours, and then 2.5 mM thymidine was added again to the cells which were then incubated for 18 hours. The efficiency of synchronization was tested by flow cytometry using cell cycle analysis kit (Multisciences).

HeLa cells were irradiated by gamma-ray provided by a Biobeam Cs137 irradiator (cat no. GM 2000; Gamma-Service Medical, Leipzig, Germany), with the doses of 1, 2, and 4 Gy at the dose rate of 3.37 Gy/ min. In the control group, the cells were taken out of the incubator without irradiation.

#### Cell viability, LDH measurements and colony-formation assays

Cell viability was measured with CCK-8 assay kit (Dojindo, Japan). Briefly, 24 h prior to irradiation treatment, ca. 6000 cells were plated in 96-well plates in a final volume of 100 μl, and then cultured for 24 h. And after adding the CCK-8 solution, the plate was incubated for another 1 h. The optical density was measured at 450 nm using a microplate reader (SpectraMax M5, Molecular Devices, USA). The amount of LDH released into the medium was assayed using the LDH-Cytotoxicity Assay Kit II (Dojindo, Japan) according to the manufacturer’s instructions. For the colony formation assay, 500 cells were seeded in 6-well plate. After being irradiated, the cells were cultured at 37℃ for about 2 weeks. After carefully washing twice with PBS, 4% paraformaldehyde was added to fix the cells. The the cells were then stained with crystal violet solution and evaluated.

#### Analysis of Cytosolic ROS and Mitochondrial ROS

Cytosolic ROS and mitochondrial ROS were detected by CellROX Green and MitoSOX Red probes, respectively. The HeLa cells were plated in corning 96-well plate at a density of 8000 cells/ well, cultivated overnight. At 6 h after irradiation, cells were incubated with 5 μM CellROX Green probe or 5 mM MitoSOX Red probe staining solution for 30 min. Then, the HeLa cells were washed with HBSS buffer thrice and incubated with 2 μg/ml Hoechst 33342 for 10 min. Fluorescent signals were analyzed by Thermo CX5 HCS fluorescence microscopy.

#### Analysis of mitochondrial membrane potential

Mitochondrial membrane potential (ΔΨ_m_) was measured using the JC-1 probe with the mitochondrial membrane potential assay kit (Beyotime, China) according to the manufacturer’s instructions. Briefly, the HeLa cells were seeded in 96 well plates with 8000 cells per well and cultured overnight at 37 ℃. After 12 hours of irradiation, the HeLa cells were incubated with 1 mg/ml JC-1 and 2 μg/ml Hoechst 33342 in dark for 20 minutes, washed with HBSS buffer for 3 times, and then photographed with Thermo CX5 HCS fluorescence microscope or quantitatively analyzed with flow cytometry. The monomer fluorescence (green) of JC-1 was observed at emission Em=521 nm with excitation Ex=485 nm, and the aggregated fluorescence (red) of JC-1 was observed at Em=607 nm with Ex=560 nm.

#### Annexin V-FITC/PI staining and cell cycle analysis

The HeLa cells were seeded at 2 × 10^5^ cell /well in 6-well plates and irradiation with gamma-ray. After 24 h, the cells were fixed with cold ethanol and then stained with propidium iodide according to kit manuscript (MultiSciences). The cells (about 20,000) were analyzed by flow cytometry (Beckman CytoFLEX, USA). The total cells were washed once and resuspended in Annexin V binding buffer, and then stained with 5 μl Annexin V and 5μl PI (Roche, Germany) for 20 min and analyzed with Beckman CytoFLEX flow cytometry. Ten thousand events were collected for each sample.

#### Evaluation of citrate synthase (CS) activity

Citrate synthase (CS) activity was determined using a citrate synthase activity assay kit (Solarbio, China). According to the instruction of the kit, the protein concentration was determined after cell lysis. Then, 7 μL protein was mixed with other reagents in the kit and put into a slit quartz cuvette. The initial absorbance A1 was recorded at 412 nm for 10 seconds. The whole quartz cuvette was then put into a 37 ℃ water bath. After 2 minutes of reaction, it was taken out, and the absorbance A2 was recorded at 412 nm, so the OD value was calculated Δ A=A2-A1. CS enzyme activity was calculated according to the formula provided with the assay kit.

#### Intracellular Ca^2+^ Measurement

The HeLa cells were loaded with the Ca^2+^ indicator Fluo-4 AM probe to assess the intracellular calcium signaling after cell irradiation at 6 h. Rhod-2 AM was used to monitor the changes in mitochondrial calcium concentrations. Fluorescent signals were analyzed by Thermo CX5 HCS fluorescence microscopy.

#### Mito-Keima mitophagy analysis

The HeLa cells were transfected with the mKeima-Red-Mito-7 plasmid using Lipo2000. After irradiation with gamma-ray, the nuclei were stained with Hoechst 33342. The fluorescence was measured with Thermo CX5 HCS fluorescence microscope and the ratio of fluorescence at 560 nm over fluorescence at 485 nm Keima fluorescence was calculated to reflect the level of mitophagy.

#### Determination of ATP levels

Intracellular ATP measured using a luciferin/luciferase assay kit (Invitrogen, USA). Cells were cultured in a black 96 well plates at density of 1 × 10^5^ cells per well. After irradiated with gamma-ray, the cells were washed with HBSS and then lysed according to the manufacturer’s instructions. Luminescence (which is proportional to the amount of ATP) was measured with SpectraMax M5 microplate reader.

#### Measurement of mitochondrial to nuclear DNA ratio (mtDNA/ nDNA)

Total genomic DNA of HeLa cells was extracted after irradiation using Total genomic DNA extract kit (TAKARA, China). Human nuclear GAPDH gene (forward: 5′-CAGAACATCATCCCTGCCTCTAC-3′; reverse: 5′-AAGGGTTGTAGTAGCCCGTAG-3′) and the mtDNA ND1 gene (forward: 5′-TTCTAATCGCAATGGCATTCCT-3′; reverse: 5′-AAGGGTTGTAGTAGCCCGTAG-3′), as well as a primer were used to detect the mtDNA copy number. The real-time quantitative PCR analysis was performed using SYBR Premix Ex Taq II (TAKARA, China) on LightCycler 480 PCR System (Roche, Germany). The calculations were analyzed with 2^-ΔΔCt^ method.

#### Analysis of mean fluorescence intensity (MFI) of dual fluorescent protein

MFI of the dual fluorescent protein was analyzed using a flow cytometer. The cells were treated with different radiation doses and collected at different time points, washed with PBS buffer, then assessed by flow cytometry (Beckman CytoFLEX, USA). The data were analyzed using Flow Jo software. The ratio of the MFI of EGFP to MFI of mecherry was used for calculation and analysis. Each irradiated dose sample group was analyzed for ca. 10,000 cells, and three measurements were repeated.

Furthermore, the High-Content Screening system Cell Insight CX5 HCS (Thermo Fisher, Waltham, MA) was used for automatic photo quantitative analysis(Sun et al., 2016). The dual-fluorescent protein HeLa cells were seeded in 96-well plates by irradiated 1 Gy, 2 Gy, 4 Gy gamma-ray. The nuclei were stained with Hoechst 33342 (Targetmol). Fluorescence ratios of protein expression of mitochondrial expressed COX8-EGFP and actin-mCherry were statistically analyzed by cellomics software (Thermo Fisher, Waltham, MA).

#### Laser scanning confocal fluorescence microscopy

All the images were obtained using 488 nm and 561 nm lasers to excite cox8-EGFP-actin-mCherry, while the 405 nm laser lines was used to excite DAPI. The imaging was performed in line mode, the z-stack was over-sampled and photographed at the height of 0.7 microns, and the image was acquired on the LEICA SP8 confocal microscope system. The contrast and brightness of the image were adjusted by Application Suite X (LEICA).

#### Western blotting analysis and immunofluorescence staining assay

Total proteins were extracted from cells using RIPA lysis buffer with a protease inhibitor cocktail (Roche, Germany). The proteins were quantified using Thermo Scientific Pierce BCA Protein Assay. The protein level of Bax, Bcl-2, caspase-3, cleaved caspase-9, Cytoc, p-AMPK, PARP, TFAM, Mfn1, PGC-1α, GFP and loading control β-Actin were determined by Western blotting assay.

For immunofluorescence staining, HeLa cells were grown on chamber slides. After γ-ray irradiation or mock irradiation, they were fixed with cold methanol for 10 minutes and permeated with 0.5% Triton X-100 in PBS for 20 minutes. After blocking for 1 hour, the cells were incubated with the primary antibody overnight and then incubated with the fluorescently labeled secondary antibody for 1 hour at 37°C. Samples investigated by laser confocal microscopy (LEICA). Image auto-quantification and co-localized foci were recognized by CellProfiler version 4 (www.cellprofiler.org)(McQuin www.cellprofiler.org)(McQuin et al., 2018).

## Statistical analysis

Data were presented as mean ± standard deviation (SD). Statistical significance was assessed by one-way or two-way ANOVA via GraphPad Prism software (Prism version 9, San Diego, CA, USA). *P*-Value < 0.05 was considered as statistically significant.

## ACKNOWLEDGEMENTS

This work was supported by the National Natural Science Foundation of China (grants No. 11635013 and No. 11775272) and Special Repair and Purchase Fund for Central-level scientific institutions (No. Y79XG13361).

## AUTHOR CONTRIBUTIONS

QH conceived the idea and supervised the research. CS and QH designed and performed the experiments, analyzed and interpreted data, performed statistical analysis, and wrote the manuscript. XZ, YM, PW and QZ provided technical assistance in the experiments. All authors read and approved the final manuscript.

## DISCLOSURES

The authors declare that they have no competing interests.

## Supplementary Information

**Figure S1.**
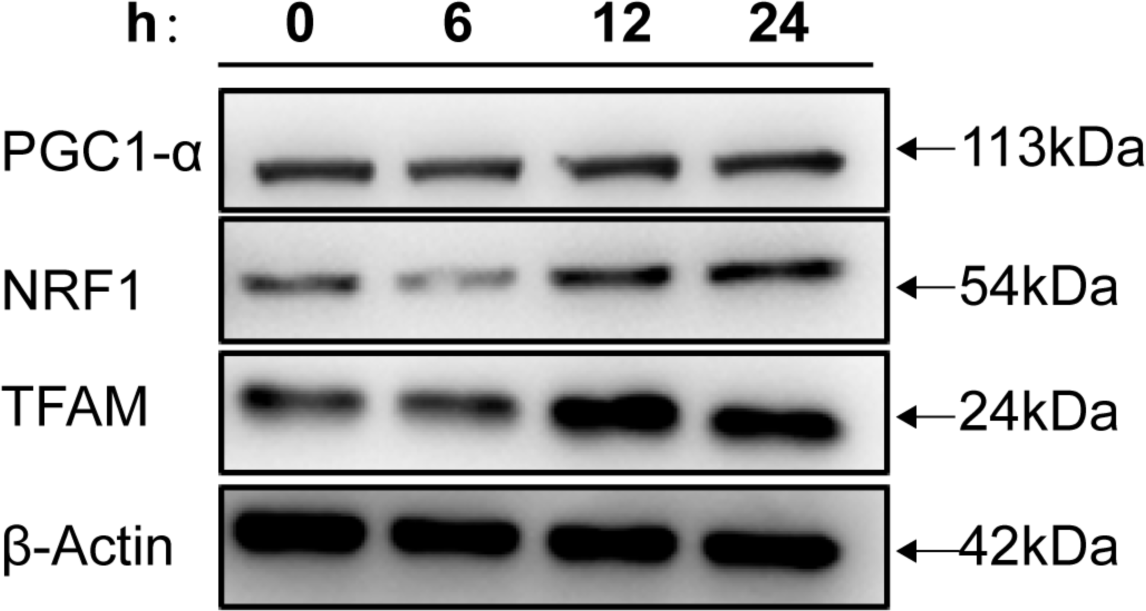
Western blot assay for verifying the effect of AICAR on mitochondrial biogenesis HeLa cells were treated with mitochondrial biogenesis inducer drug (AICAR, 0.5 mM), and the whole cell extracts were collected at different time points. The mitochondrial biogenesis marker proteins (PGC-1α, NRF1 and TFAM) are indicated in the plot.

**Figure S2.**
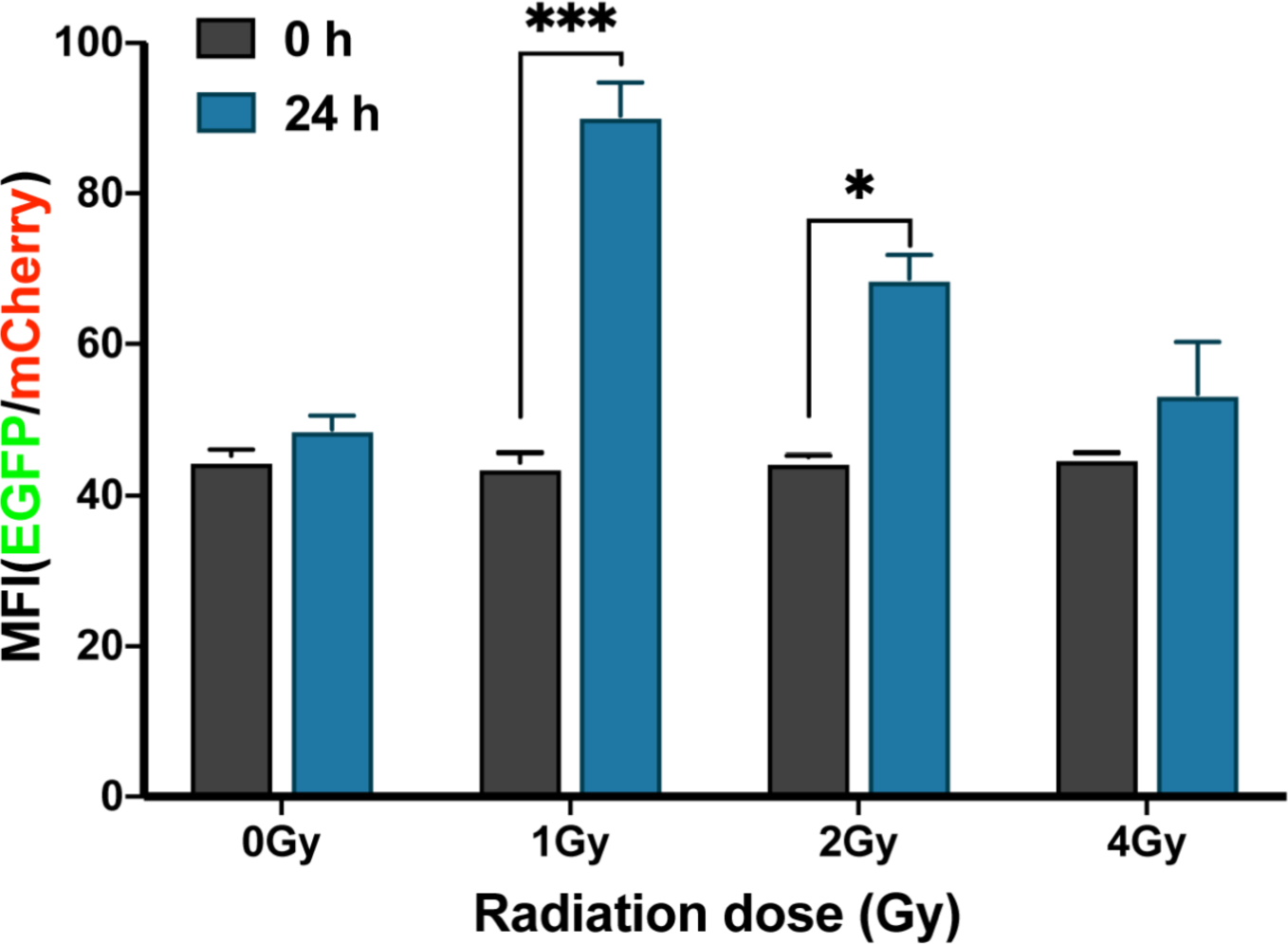
The mean fluorescence intensity (MFI) was determined by flow cytometry for the HeLa cells with or without irradiation. HeLa-cox8-EGFP-actin-mCherry cells were exposed to 1 Gy, 2 Gy, 4 Gy gamma-ray and incubated for the indicated periods. The analysis and comparison MFI obtained from the dual-fluorescent measurements by flow cytometry.

**Figure S3.**
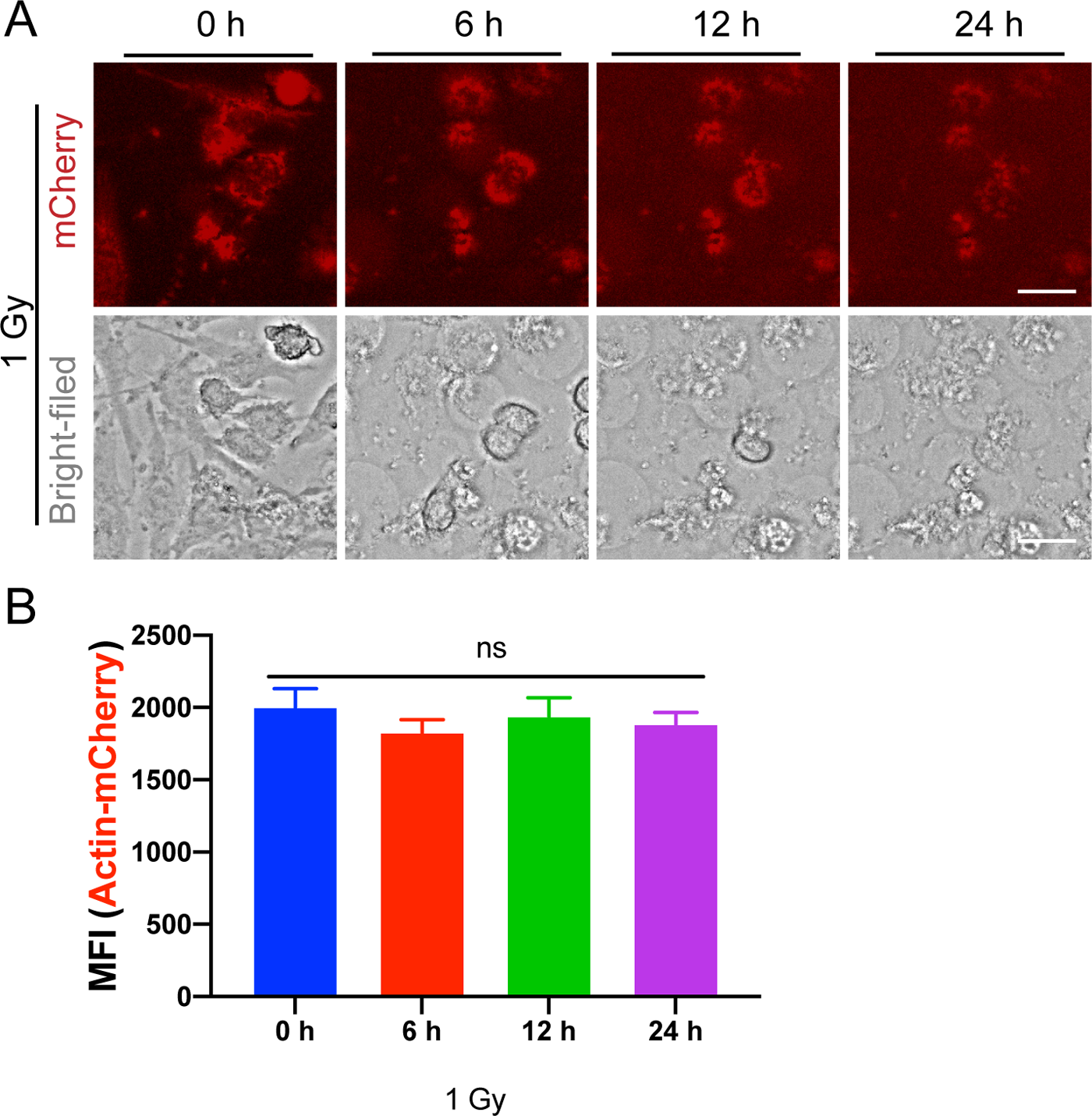
The mean fluorescence intensity (MFI) of Actin-mCherry was determined by living fluorescence microscopy for the HeLa cells with with or without irradiation (1 Gy). (A) Fluorescence images and bright field images were taken at 0, 6, 12 and 24 hours after irradiatied with 1 Gy dose. (B) The histogram is the statistical chart of the mean fluorescence intensity of Actin-mCherry.

**Figure S4.**
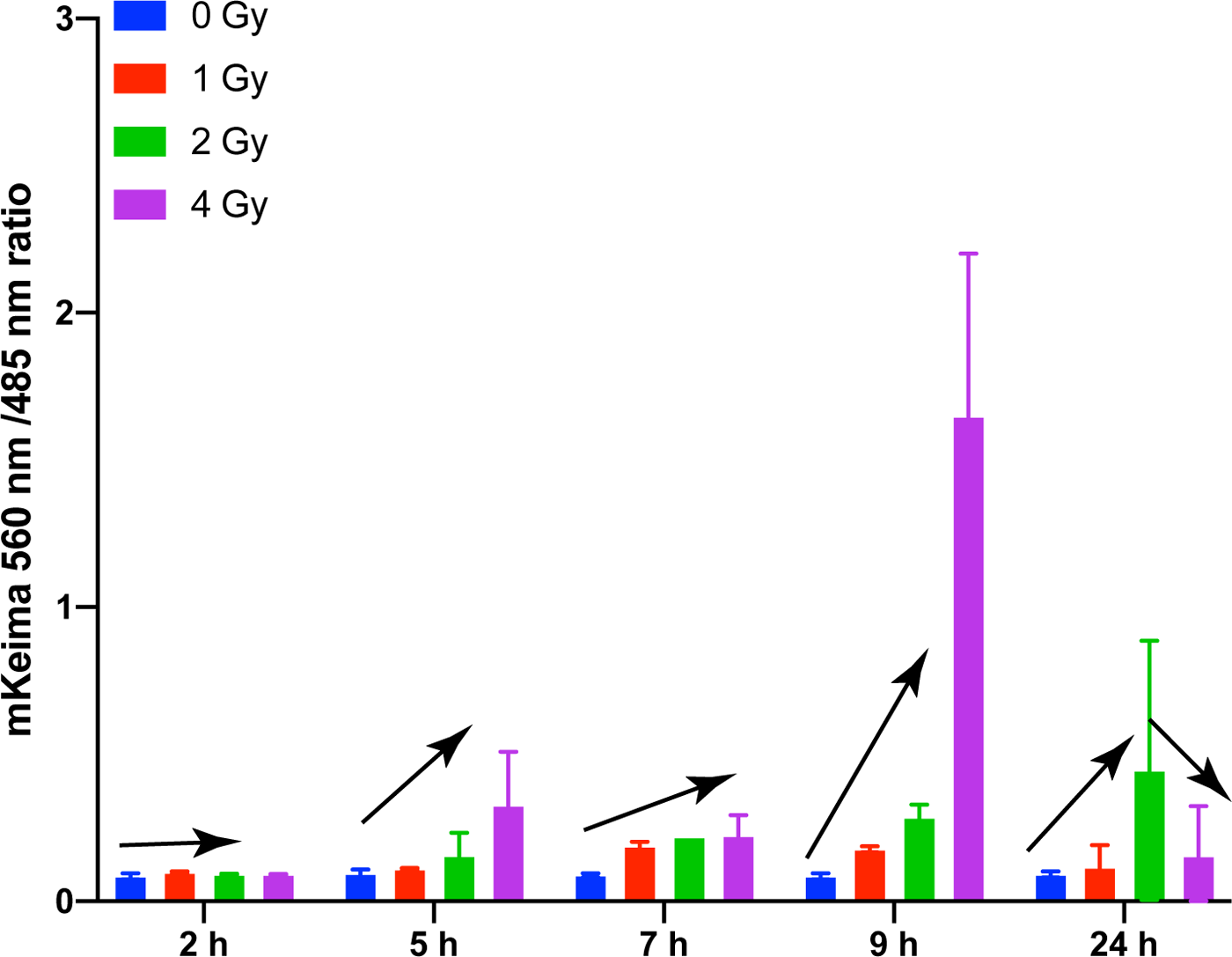
Mitophagy activity assessed using mt-Keima in HeLa cells Elevated levels of mitophagy were observed following irradiation with 1 Gy, 2 Gy, 4 Gy. The fluorescence ratio of red fluorescence to green fluorescence (560 nm/ 485 nm) at different time points (2, 5, 7, 9 and 24 hours) was calculated by high content fluorescence analysis system. n = 1000 cells calculated per group.

**Table S1.**
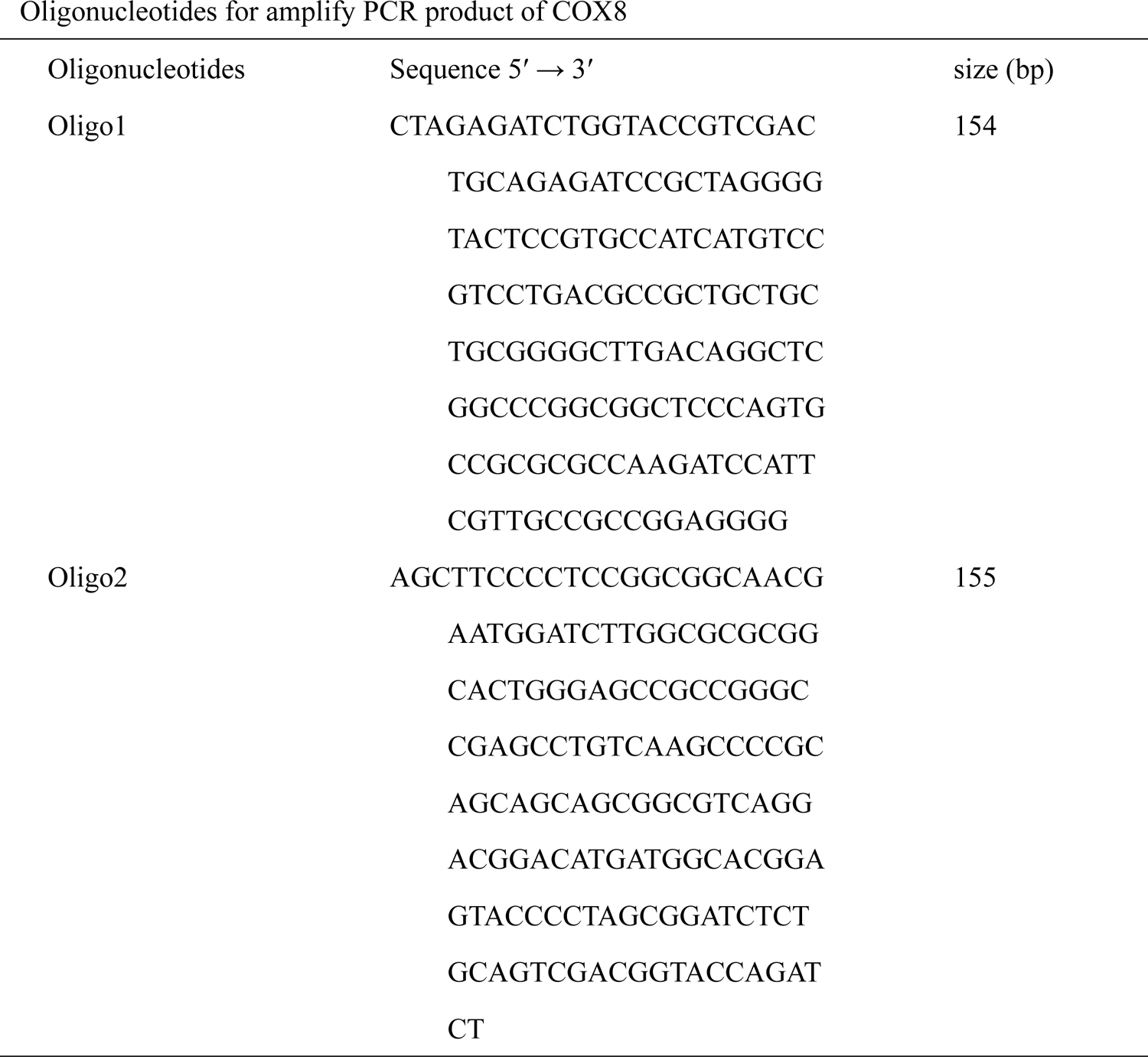
Oligonucleotides for amplify PCR product of COX8

---

**S1 Video**

**S2 Video**

**S3 Video**

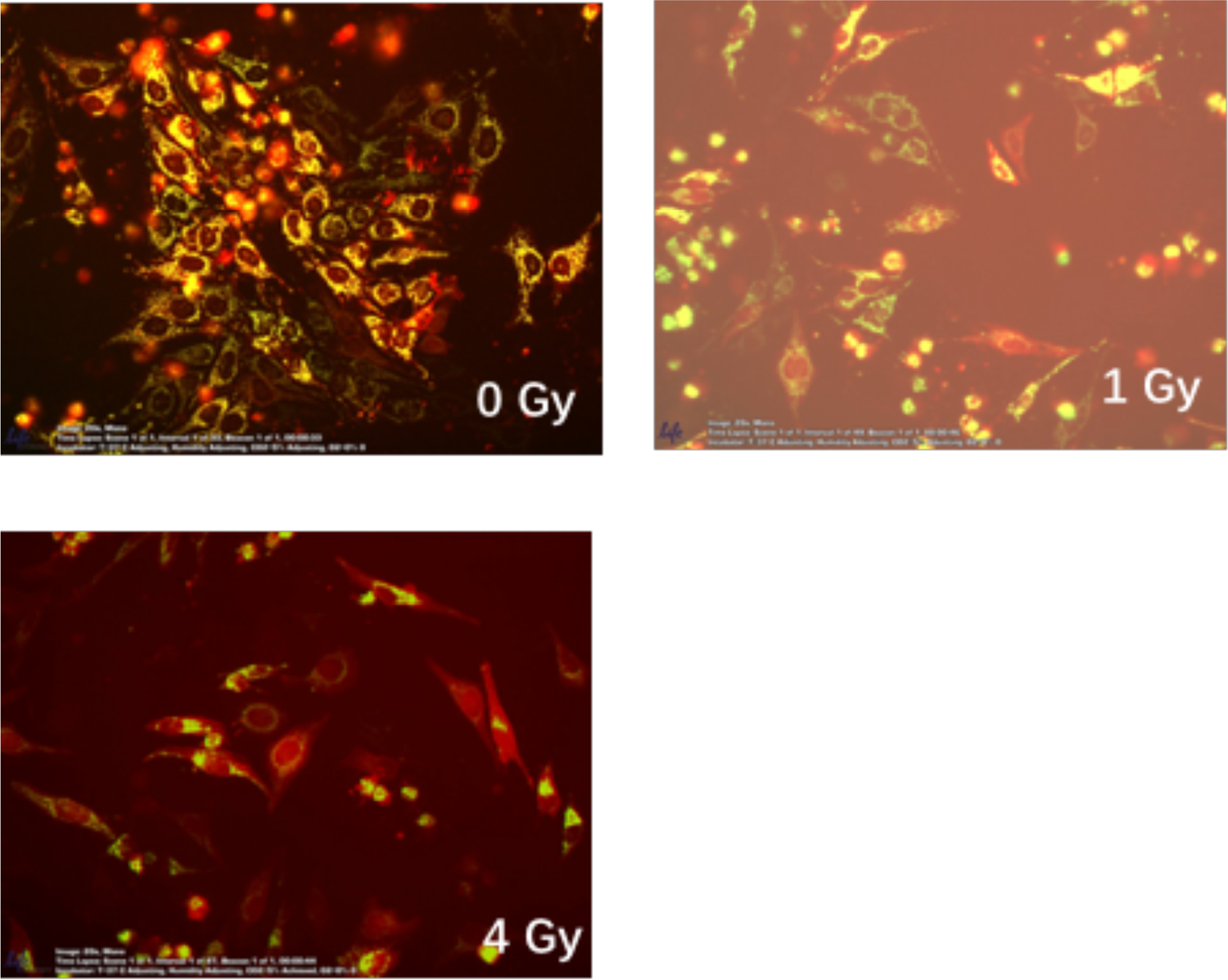

In-situ real-time video of dual fluorescence reporter system of irradiated cells (Video S1, Video S2, Video S3)

## Notes

### Competing Interest Statement

The authors have declared no competing interest.

## References

1. Azzam, E.I., Jay-Gerin, J.-P., and Pain, D. (2012). Ionizing radiation-induced metabolic oxidative stress and prolonged cell injury. Cancer Lett 327, 48–60.

2. Barros, M.H., and McStay, G.P. (2019). Modular Biogenesis of Mitochondrial Respiratory Complexes. Mitochondrion 50, 94–114.

3. Bernstein, C., Bernstein, H., Payne, C.M., and Garewal, H. (2002). DNA repair/pro-apoptotic dual-role proteins in five major DNA repair pathways: fail-safe protection against carcinogenesis. Mutat Res Rev Mutat Res 511, 145–178.

4. Betzig, E., Patterson, G.H., Sougrat, R., Lindwasser, O.W., Olenych, S., Bonifacino, J.S., Davidson, M.W., Lippincott-Schwartz, J., and Hess, H.F. (2006). Imaging Intracellular Fluorescent Proteins at Nanometer Resolution. Science 313, 1642–1645.

5. Bost, F., and Kaminski, L. (2019). The metabolic modulator PGC-1α in cancer. Am J Cancer Res 9, 198–211.

6. Burke, P.J. (2017). Mitochondria, Bioenergetics and Apoptosis in Cancer. Trends Cancer 3, 857– 870.

7. Cao, C., Yu, H., Wu, F., Qi, H., and He, J. (2017). Antibiotic anisomycin induces cell cycle arrest and apoptosis through inhibiting mitochondrial biogenesis in osteosarcoma. J Bioenerg Biomembr 49, 437–443.

8. Chalkiadaki, A., and Guarente, L. (2015). The multifaceted functions of sirtuins in cancer. Nat Rev Cancer 15, 608–624.

9. Dam, A.D., Mitchell, A.S., and Quadrilatero, J. (2013). Induction of mitochondrial biogenesis protects against caspase-dependent and caspase-independent apoptosis in L6 myoblasts. Biochimica Et Biophysica Acta Bba - Mol Cell Res 1833, 3426–3435.

10. Das, S., Joshi, M.B., Parashiva, G.K., and Rao, S.B.S. (2020). Stimulation of cytoprotective autophagy and components of mitochondrial biogenesis / proteostasis in response to ionizing radiation as a credible pro-survival strategy. Free Radical Bio Med.

11. Davis, A.J., Chen, B.P.C., and Chen, D.J. (2014). DNA-PK: A dynamic enzyme in a versatile DSB repair pathway. Dna Repair 17, 21–29.

12. Fanibunda, S.E., Deb, S., Maniyadath, B., Tiwari, P., Ghai, U., Gupta, S., Figueiredo, D., Weisstaub, N., Gingrich, J.A., Vaidya, A.D.B., et al. (2019). Serotonin regulates mitochondrial biogenesis and function in rodent cortical neurons via the 5-HT2A receptor and SIRT1–PGC-1α axis. Proc National Acad Sci 116, 201821332.

13. Fu, X., Wan, S., Lyu, Y.L., Liu, L.F., and Qi, H. (2008). Etoposide Induces ATM-Dependent Mitochondrial Biogenesis through AMPK Activation. Plos One 3, e2009.

14. Galluzzi, L., and Kroemer, G. (2008). Necroptosis: a specialized pathway of programmed necrosis. Cell 135, 1161–1163.

15. Haggie, P.M., and Verkman, A.S. (2002). Diffusion of Tricarboxylic Acid Cycle Enzymes in the Mitochondrial Matrixin Vivo: EVIDENCE FOR RESTRICTED MOBILITY OF A MULTIENZYME COMPLEX. J Biol Chem 277, 40782–40788.

16. Hanson, G.T., Aggeler, R., Oglesbee, D., Cannon, M., Capaldi, R.A., Tsien, R.Y., and Remington, S.J. (2004). Investigating Mitochondrial Redox Potential with Redox-sensitive Green Fluorescent Protein Indicators. J Biol Chem 279, 13044–13053.

17. Holler, N., Zaru, R., Micheau, O., Thome, M., Attinger, A., Valitutti, S., Bodmer, J.-L., Schneider, P., Seed, B., and Tschopp, J. (2000). Fas triggers an alternative, caspase-8–independent cell death pathway using the kinase RIP as effector molecule. Nat Immunol 1, 489–495.

18. Huangyang, P., Li, F., Lee, P., Nissim, I., Weljie, A.M., Mancuso, A., Li, B., Keith, B., Yoon, S.S., and Simon, M.C. (2020). Fructose-1,6-Bisphosphatase 2 Inhibits Sarcoma Progression by Restraining Mitochondrial Biogenesis. Cell Metab 31, 1032.

19. Jadhav, U., and Shivdasani, R.A. (2019). Dissecting Cell Lineages: From Microscope to Kaleidoscope. Cell 176, 949–951.

20. Katajisto, P., Döhla, J., Chaffer, C.L., Pentinmikko, N., Marjanovic, N., Iqbal, S., Zoncu, R., Chen, W., Weinberg, R.A., and Sabatini, D.M. (2015). Stem cells. Asymmetric apportioning of aged mitochondria between daughter cells is required for stemness. Sci New York N Y 348, 340– 343.

21. Kim, J., Yang, G., Kim, Y., Kim, J., and Ha, J. (2016). AMPK activators: mechanisms of action and physiological activities. Exp Mol Medicine 48, e224–e224.

22. Kluza, J., Marchetti, P., Gallego, M.-A., Lancel, S., Fournier, C., Loyens, A., Beauvillain, J.-C., and Bailly, C. (2004). Mitochondrial proliferation during apoptosis induced by anticancer agents: effects of doxorubicin and mitoxantrone on cancer and cardiac cells. Oncogene 23, 7018–7030.

23. Komen, J.C., and Thorburn, D.R. (2014). Turn up the power – pharmacological activation of mitochondrial biogenesis in mouse models. Brit J Pharmacol 171, 1818–1836.

24. Kulms, D., Zeise, E., Pöppelmann, B., and Schwarz, T. (2002). DNA damage, death receptor activation and reactive oxygen species contribute to ultraviolet radiation-induced apoptosis in an essential and independent way. Oncogene 21, 5844–5851.

25. Leach, J.K., Tuyle, G.V., Lin, P.S., Schmidt-Ullrich, R., and Mikkelsen, R.B. (2001). Ionizing radiation-induced, mitochondria-dependent generation of reactive oxygen/nitrogen. Cancer Res 61, 3894–3901.

26. Lee, H.-C., and Wei, Y.-H. (2005). Mitochondrial biogenesis and mitochondrial DNA maintenance of mammalian cells under oxidative stress. Int J Biochem Cell Biology 37, 822–834.

27. Li, S., Shi, B., Liu, X., and An, H.-X. (2020). Acetylation and Deacetylation of DNA Repair Proteins in Cancers. Frontiers Oncol 10, 573502.

28. Lill, R., and Freibert, S.-A. (2020). Mechanisms of Mitochondrial Iron-Sulfur Protein Biogenesis. Annu Rev Biochem 89, 471–499.

29. Lin, W.T., Nithiyanantham, S., Hsieh, D.J., Chen, R., Day, C., Liao, J.Y., Kuo, C., Mahalakshmi, B., Kuo, W., and Huang, C. (2020). Bioactive peptides attenuate cardiac apoptosis in spontaneously hypertensive rat hearts through activation of autophagy and mitochondrial biogenesis pathway. Environ Toxicol 35, 804–810.

30. Llopis, J., McCaffery, J.M., Miyawaki, A., Farquhar, M.G., and Tsien, R.Y. (1998). Measurement of cytosolic, mitochondrial, and Golgi pH in single living cells with green fluorescent proteins. Proc National Acad Sci 95, 6803–6808.

31. Löbrich, M., and Jeggo, P.A. (2005). Harmonising the response to DSBs: a new string in the ATM bow. Dna Repair 4, 749–759.

32. Mahajan, N.P., Linder, K., Berry, G., Gordon, G.W., Heim, R., and Herman, B. (1998). Bcl-2 and Bax interactions in mitochondria probed with green fluorescent protein and fluorescence resonance energy transfer. Nat Biotechnol 16, 547–552.

33. Martins, L.A.M., Vieira, M.Q., Ilha, M., Vasconcelos, M. de Biehl, H.B., Lima, D.B., Schein, V., Barbé-Tuana, F., Borojevic, R., and Guma, F.C.R. (2015). The Interplay Between Apoptosis, Mitophagy and Mitochondrial Biogenesis Induced by Resveratrol Can Determine Activated Hepatic Stellate Cells Death or Survival. Cell Biochem Biophys 71, 657–672.

34. May-Panloup, P., Vignon, X., Chrétien, M.-F., Heyman, Y., Tamassia, M., Malthièry, Y., and Reynier, P. (2005). Increase of mitochondrial DNA content and transcripts in early bovine embryogenesis associated with upregulation of mtTFA and NRF1 transcription factors. Reprod Biol Endocrin 3, 65.

35. McQuin, C., Goodman, A., Chernyshev, V., Kamentsky, L., Cimini, B.A., Karhohs, K.W., Doan, M., Ding, L., Rafelski, S.M., Thirstrup, D., et al. (2018). CellProfiler 3.0: Next-generation image processing for biology. Plos Biol 16, e2005970.

36. Melentijevic, I., Toth, M.L., Arnold, M.L., Guasp, R.J., Harinath, G., Nguyen, K.C., Taub, D., Parker, J.A., Neri, C., Gabel, C.V., et al. (2017). C. elegans neurons jettison protein aggregates and mitochondria under neurotoxic stress. Nature 542, 367–371.

37. Mihaylova, M.M., and Shaw, R.J. (2011). The AMPK signalling pathway coordinates cell growth, autophagy and metabolism. Nat Cell Biol 13, 1016–1023.

38. Miyashiro, T., and Goulian, M. (2007). Methods in Enzymology. Methods Enzymol 423, 458– 475.

39. Montava-Garriga, L., and Ganley, I.G. (2019). Outstanding Questions in Mitophagy: What we Do and Do Not Know. J Mol Biol 432, 206–230.

40. Mukhopadhyay, P., Rajesh, M., Haskó, G., Hawkins, B.J., Madesh, M., and Pacher, P. (2007). Simultaneous detection of apoptosis and mitochondrial superoxide production in live cells by flow cytometry and confocal microscopy. Nat Protoc 2, 2295–2301.

41. Newman, R.H., Fosbrink, M.D., and Zhang, J. (2011). Genetically encodable fluorescent biosensors for tracking signaling dynamics in living cells. Chem Rev 111, 3614–3666.

42. O’Driscoll, M., and Jeggo, P.A. (2006). The role of double-strand break repair — insights from human genetics. Nat Rev Genet 7, 45–54.

43. Paredes, S., and Chua, K.F. (2016). SIRT7 clears the way for DNA repair. Embo J 35, 1483–1485.

44. Park, S.-J., Gavrilova, O., Brown, A.L., Soto, J.E., Bremner, S., Kim, J., Xu, X., Yang, S., Um, J.-H., Koch, L.G., et al. (2017). DNA-PK Promotes the Mitochondrial, Metabolic, and Physical Decline that Occurs During Aging. Cell Metab 25, 1135–1146.e7.

45. Popov, L. (2020). Mitochondrial biogenesis: An update. J Cell Mol Med 24, 4892–4899.

46. Rai, P.K., Russell, O.M., Lightowlers, R.N., and Turnbull, D.M. (2015). Potential compounds for the treatment of mitochondrial disease. Brit Med Bull 116, 5–18.

47. Rai, Y., Pathak, R., Kumari, N., Sah, D.K., Pandey, S., Kalra, N., Soni, R., Dwarakanath, B.S., and Bhatt, A.N. (2018). Mitochondrial biogenesis and metabolic hyperactivation limits the application of MTT assay in the estimation of radiation induced growth inhibition. Sci Rep-Uk 8, 1531.

48. Ruan, L., Zhou, C., Jin, E., Kucharavy, A., Zhang, Y., Wen, Z., Florens, L., and Li, R. (2017). Cytosolic proteostasis through importing of misfolded proteins into mitochondria. Nature 543, 443–446.

49. Scarpulla, R.C. (2008). Transcriptional Paradigms in Mammalian Mitochondrial Biogenesis and Function. Physiol Rev 88, 611–638.

50. Spitz, D.R., Azzam, E.I., Li, J.J., and Gius, D. (2004). Metabolic oxidation/reduction reactions and cellular responses to ionizing radiation: A unifying concept in stress response biology. Cancer Metast Rev 23, 311–322.

51. Srinivas, U.S., Tan, B.W.Q., Vellayappan, B.A., and Jeyasekharan, A.D. (2019). ROS and the DNA damage response in cancer. Redox Biol 25, 101084.

52. Sun, J., Li, N., Oh, K.-S., Dutta, B., Vayttaden, S.J., Lin, B., Ebert, T.S., Nardo, D.D., Davis, J., Bagirzadeh, R., et al. (2016). Comprehensive RNAi-based screening of human and mouse TLR pathways identifies species-specific preferences in signaling protein use. Sci Signal 9, ra3–ra3.

53. Vayssiere, J.L., Petit, P.X., Risler, Y., and Mignotte, B. (1994). Commitment to apoptosis is associated with changes in mitochondrial biogenesis and activity in cell lines conditionally immortalized with simian virus 40. Proc National Acad Sci 91, 11752–11756.

54. Virbasius, C.A., Virbasius, J.V., and Scarpulla, R.C. (1993). NRF-1, an activator involved in nuclear-mitochondrial interactions, utilizes a new DNA-binding domain conserved in a family of developmental regulators. Gene Dev 7, 2431–2445.

55. Wang, Y., An, H., Liu, T., Qin, C., Sesaki, H., Guo, S., Radovick, S., Hussain, M., Maheshwari, A., Wondisford, F.E., et al. (2019). Metformin Improves Mitochondrial Respiratory Activity through Activation of AMPK. Cell Reports 29, 1511–1523.e5.

56. Westermann, B., and Neupert, W. (2000). Mitochondria-targeted green fluorescent proteins: convenient tools for the study of organelle biogenesis inSaccharomyces cerevisiae. Yeast 16, 1421–1427.

57. Wu, Z., Puigserver, P., Andersson, U., Zhang, C., Adelmant, G., Mootha, V., Troy, A., Cinti, S., Lowell, B., Scarpulla, R.C., et al. (1999). Mechanisms Controlling Mitochondrial Biogenesis and Respiration through the Thermogenic Coactivator PGC-1. Cell 98, 115–124.

58. Yambire, K.F., Fernandez-Mosquera, L., Steinfeld, R., Mühle, C., Ikonen, E., Milosevic, I., and Raimundo, N. (2019). Mitochondrial biogenesis is transcriptionally repressed in lysosomal lipid storage diseases. Elife 8, e39598.

59. Yu, M. (2011). Generation, function and diagnostic value of mitochondrial DNA copy number alterations in human cancers. Life Sci 89, 65–71.

60. Yu, H., Zhao, W., Xie, M., Li, X., Sun, M., He, J., Wang, L., and Yu, L. (2020). Real-Time Monitoring of Self-Aggregation of β-Amyloid by a Fluorescent Probe Based on Ruthenium Complex. Anal Chem 92, 2953–2960.

61. Yu, J., Wang, Q., Chen, N., Sun, Y., Wang, X., Wu, L., Chen, S., Yuan, H., Xu, A., and Wang, J. (2013). Mitochondrial transcription factor A regulated ionizing radiation-induced mitochondrial biogenesis in human lung adenocarcinoma A549 cells. J Radiat Res 54, 998–1004.

62. Zhang, T., Ikejima, T., Li, L., Wu, R., Yuan, X., Zhao, J., Wang, Y., and Peng, S. (2017). Impairment of Mitochondrial Biogenesis and Dynamics Involved in Isoniazid-Induced Apoptosis of HepG2 Cells Was Alleviated by p38 MAPK Pathway. Front Pharmacol 8, 753.

